# Decades of dreams coming true: capillary zone electrophoresis-mass spectrometry for reproducible multi-level proteomics

**DOI:** 10.64898/2026.01.28.702308

**Authors:** Guijie Zhu, Yifan Yue, Jorge A Colón Rosado, Guangyao Gao, Xiaowen Liu, Liangliang Sun

## Abstract

Capillary zone electrophoresis (CZE)-mass spectrometry (MS) has been proposed as a powerful analytical tool for bottom-up, top-down, and native proteomics (multi-level proteomics) decades ago to analyze complex biological samples at the levels of peptides (bottom-up), proteoforms (top-down), and complexoforms (native). However, its broad adoption has been impeded by the limited robustness and reproducibility. Here, we present multi-level proteomics data from nearly 170 CZE-MS runs (∼170 hours of instrument time), demonstrating qualitatively (i.e., the number of identified peptides and proteoforms, the number of detected complexoforms, and their migration time) and quantitatively (i.e., peptide, proteoform, and complexoform intensity) reproducible measurement of complex samples with varying levels of complexity, i.e., *Escherichia coli* cells, HeLa cells, and human plasma. CZE-MS-based native proteomics enabled the detection of hundreds of complexoforms up to 800 kDa from the complex systems via consuming only nanograms of protein material. The results indicate that CZE-MS is sensitive and reproducible enough for broad adoption for multi-level proteomics-based biomedical research.

---

According to PubMed, more than 30,000 papers on capillary electrophoresis (CE) and nearly 6,000 papers on CE-mass spectrometry (MS) were published since the 1980s, with more than 200 CE-MS papers per year in the past 20 years. CE-MS has been applied in a variety of applications, including biomedical, clinical, environmental, and food analysis. Capillary zone electrophoresis-MS (CZE-MS) has been recognized as a useful tool for multi-level proteomics, i.e., bottom-up (BUP), top-down (TDP), and native proteomics, for measuring peptides, intact proteoforms [1], and complexoforms [2] decades ago. [3-11] CZE-MS has many valuable features for multi-level proteomics, including high sensitivity [12-18], high separation efficiency [19,20], the capability of maintaining the integrity and topology of protein complexes [21-26], and accurate prediction of electrophoretic mobility of analytes [27-29]. However, it has not been widely adopted in proteomics due to issues of robustness and reproducibility. The maturity of several CE-MS interfaces [30-33] and the improvement of capillary inner-wall coatings [34] have substantially advanced the performance of CZE-MS for multi-level proteomics. [9, 35-37] However, systematic evaluations of advanced CZE-MS in terms of long-term reproducibility and robustness for multi-level proteomics are missing, impeding its broad adoption in biomedical research.

Here, for the first time, we investigated the long-term reproducibility of CZE-MS for BUP, TDP, and native proteomics using biological samples with varied sample complexities, i.e., *Escherichia coli* (*E. coli)* cells, HeLa cells, and human plasma. We employed one commercial CE-MS interface, an electrokinetically pumped sheath-flow CE-MS interface (EMASS-II, CMP Scientific), [31,38] for the experiment. Linear polyacrylamide (LPA)-coated fused silica capillaries were used for CZE separation. [34,39] The dynamic pH junction method [20, 40, 41] was used to perform online sample stacking for BUP and TDP experiments. The experimental details are described in **Supporting Information I**. We performed BUP analysis of a commercial HeLa cell digest for 67 runs with a 70-minute instrument time per run. We conducted TDP analysis of an *E. coli* cell lysate for 26 runs and a HeLa cell lysate for approximately 30 runs. CZE-MS/MS produced reproducible BUP and TDP measurements of these complex samples. **Figure 1A** shows the example electropherograms of the BUP and TDP measurements with consistent separation profiles. **Figures S1-S3** show more examples of the reproducible electropherograms. BUP analysis of the HeLa cell digest across 67 CZE-MS/MS runs generated reproducible numbers of peptides, protein groups, and peptide-spectrum-matches (PSMs) with relative standard deviations (RSDs) smaller than 9%, **Figure S4A**. TDP analysis of an *E. coli* cell lysate across 26 CZE-MS/MS runs identified consistent numbers of proteoforms, proteins, and proteoform-spectrum matches (PrSMs) with RSDs less than 10%, **Figure S4A**. The number of proteoforms, proteins, and PrSMs identified from the HeLa cell lysate across 29 CZE-MS/MS runs is also reasonably consistent, but with larger RSDs than that of the *E. coli* data (∼20% vs. <10%). The larger variations in the number of proteoform identifications for the HeLa cell lysate are most likely due to the substantially higher sample complexity of HeLa cells compared to *E. coli* cells. All the identified proteoforms from the *E. coli* and HeLa cell lysates are about 30 kDa or smaller, **Figure S4B**, due to the challenges of TDP for measuring large proteoforms. [42,43]

**Figure 1.**
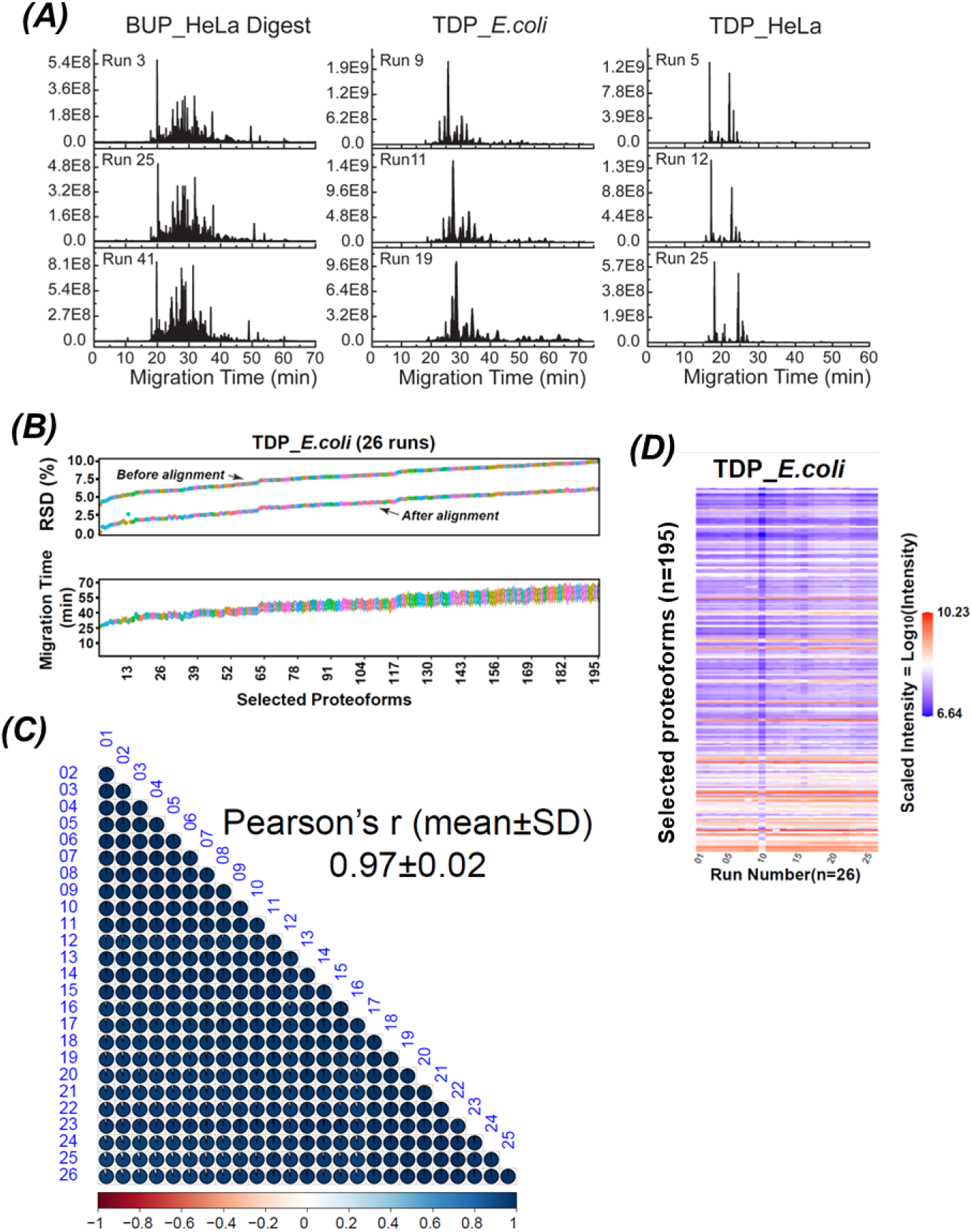
Summary of the reproducibility data of CZE-MS-based bottom-up proteomics (BUP) and top-down proteomics (TDP). (A) Example electropherograms of BUP data (HeLa cell lysate digest) and TDP data (E. coli and HeLa cell lysates). (B) Box-plots and relative standard deviations (RSDs) of migration time of selected 195 proteoforms across 26 CZE-MS runs of an E. coli cell lysate. For the RSD plots, one plot was from the original migration time data of proteoforms (before alignment), and the other one was from the migration time data after alignment based on one proteoform across the 26 runs (after alignment). (C) Pair-wise linear correlations of proteoform intensity from CZE-MS analyses of an E. coli cell lysate. The color bar shows the range of Pearson’+s r of the linear correlations. SD: standard deviation. (D) Heatmap of log10-scaled intensity of 195 selected proteoforms (y-axis) across 26 CZE-MS runs (x-axis) of an E. coli cell lysate.

We then further investigated the reproducibility of BUP and TDP regarding the migration time of peptides and proteoforms. The RSDs of migration time of 400 selected peptides are smaller than 10% across the 67 CZE-MS/MS runs, with larger RSDs of migration time for slow-migrating peptides, **Figure S5**. The slight migration time shift is due to the slight lab temperature change and minor changes in the capillary inner wall coating during the long-term evaluation. Using a simple migration time alignment approach across the 67 runs based on a single peptide, the RSDs of those peptides can be reduced to approximately less than 7%, **Figure S5**. For the TDP analysis of *E. coli* and HeLa cell lysates, CZE-MS produced reasonably consistent migration time across nearly 30 runs, with the RSDs of migration time of selected proteoforms smaller than 10%, **Figure 1B** and **Figure S6**. Similar to the BUP data, after a simple migration time alignment using one proteoform, the migration time RSDs were reduced to less than 7%. The results clearly indicate that CZE-MS can produce reproducible separation and identification of peptides and proteoforms in complex biological samples.

We then studied the reproducibility of BUP and TDP measurements by CZE-MS, focusing on peptide intensity and proteoform intensity. As shown in **Figure S7**, the pairwise linear correlation of peptide intensity from BUP analysis shows an excellent Pearson’+s *r* of 0.94±0.04 (mean±SD). **Figure 1C** shows the pairwise linear correlation of *E. coli* proteoform intensity with Pearson’+s *r* of 0.97± 0.02 (mean±SD), suggesting nice linear correlations of proteoform intensity between any two CZE-MS/MS runs. **Figure 1D** shows the heat map of the intensity of selected 195 *E. coli* proteoforms across the 26 CZE-MS/MS runs. The proteoforms covering a three-orders-of-magnitude intensity range show reasonably consistent intensity profiles across the CZE-MS/MS runs. We then analyzed the HeLa proteoform intensity data from 29 CZE-MS/MS runs by performing the pairwise linear correlations of proteoform intensity, **Figure S8**. CZE-MS/MS analysis of the HeLa cell lysate produced nice linear correlations of proteoform intensity between any two runs, with Pearson’+s *r* of 0.88±0.04 (mean±SD). The results here documented the nice reproducibility of CZE-MS/MS for BUP and TDP of complex samples regarding proteoform intensity, suggesting that CZE-MS/MS is ready for large-scale quantitative BUP and TDP studies.

Recently, native CZE-MS has been recognized as a powerful tool for native proteomics, measuring endogenous complexoforms in complex samples. [21-26] Here, for the first time, we studied the reproducibility of CZE-MS-based native proteomics for measuring biological samples with various complexities, i.e., *E. coli* cells (∼10 runs), HeLa cells (∼20 runs), and human plasma (∼20 runs), to facilitate the broad adoption of CZE-MS for native proteomics applications. **Figure 2** shows the example electropherograms of CZE-MS for native proteomics analysis of *E. coli* cells, HeLa cells, and human plasma (A), as well as the mass distributions of detected complexoforms (B). Native CZE-MS produced consistent separation profiles of complexoforms across triplicate measurements of different samples, **Figure 2A**. Native CZE-MS detected about 170 native proteoforms or complexoforms from the *E. coli* cell lysate with masses up to 800 kDa, **Figure 2B**. It measured nearly 50 native proteoforms or complexoforms from the HeLa cell lysate (i.e., one size exclusion chromatography (SEC) fraction of the whole HeLa cell lysate), and these complexoforms have masses up to 500 kDa. The native CZE-MS technique enabled the detection of over 50 native proteoforms or complexoforms from the human plasma sample with masses up to over 800 kDa. The data demonstrates the reproducibility of native CZE-MS-based native proteomics and the capability of native CZE-MS for measuring large native proteoforms or complexoforms in complex samples. We need to highlight that only roughly 350 ng or less complexoform materials were loaded for each native CZE-MS run, indicating the high sensitivity of our CZE-MS system for native proteomics applications.

**Figure 2.**
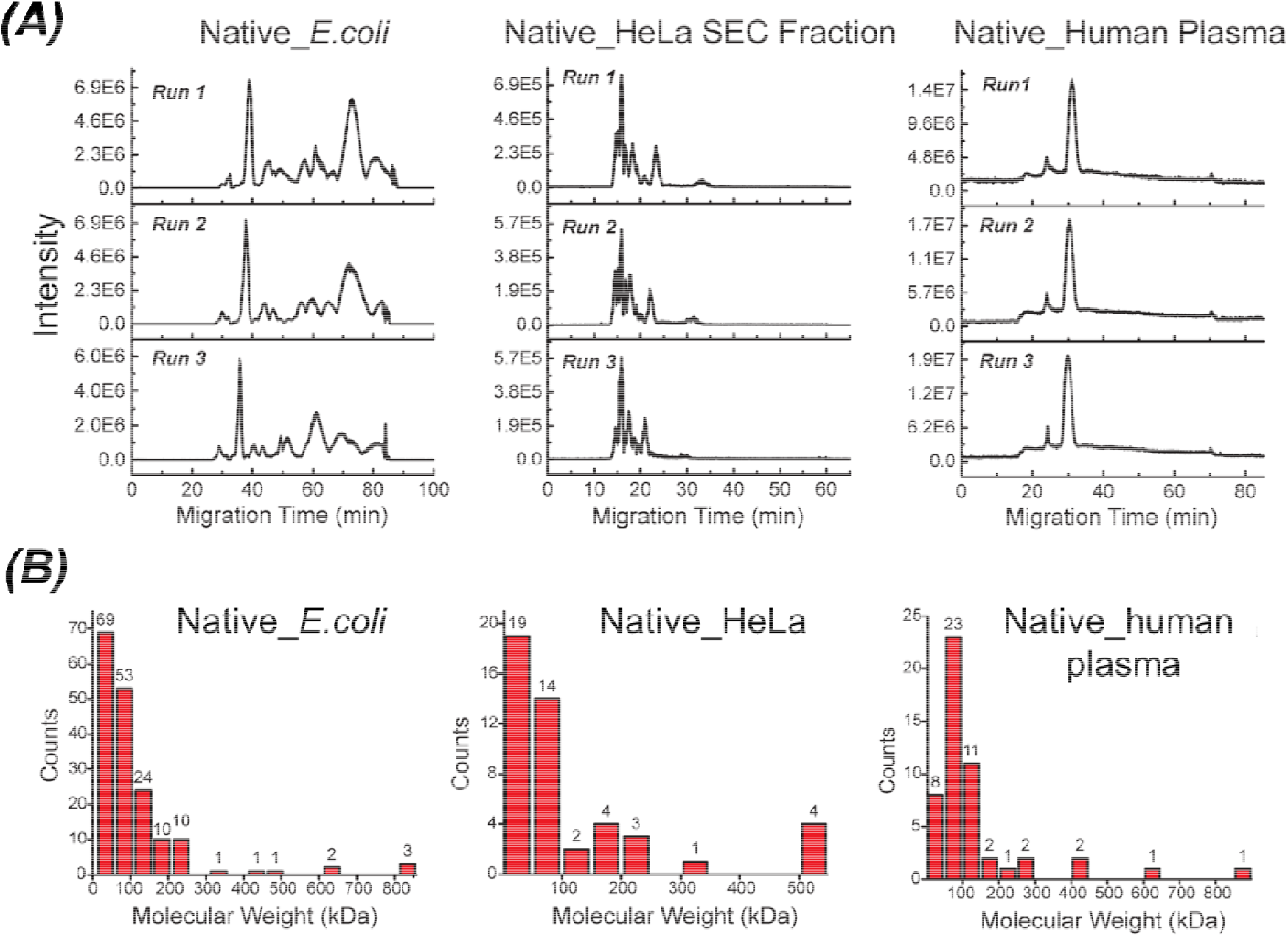
Brief summary of the native proteomics data of various biological samples. (A) Example electropherograms (triplicate runs) of CZE-MS-based native proteomics of an *E. coli* cell lysate (∼3 mg/mL), a HeLa cell lysate (one size exclusion chromatography, SEC, fraction), and human plasma. (B) Mass distributions of detected complexoforms from the three different biological samples.

**Figures S9-S19** show the electropherograms of native CZE-MS or MS/MS analyses of *E. coli* cell lysates (**Figures S9** and **S10**), one size exclusion chromatography (SEC) fraction of a HeLa cell lysate (**Figures S11-S13**), and one human plasma sample (**Figures S14-S16**). CZE-MS and MS/MS yielded reproducible measurements of complexoforms in these different samples regarding separation profiles and normalized level (NL) base peak or total ion current (TIC) intensities. The RSDs of the NL base peak or TIC intensities are less than 20% for all cases except the data in **Figure S10**, which had an RSD of 37% in NL base peak intensity, most likely due to the relatively low intensity of complexoforms. The data further highlights the nice reproducibility of native CZE-MS for different complex samples across dozens of runs.

Native CZE-MS detected hundreds of complexoforms from the three types of biological samples, i.e., *E. coli* cells, HeLa cells, and human plasma. The lists of detected complexoforms are shown in **Supporting Information II**. To identify the complexoforms, we performed CZE-MS/MS analysis of the samples in data-dependent acquisition (DDA) mode. We tried to identify the complexoforms by database search using a modified version of the TopPIC software. [44] The details of the data analysis are described in the **Supporting Information I**.

We identified multiple complexoforms by MS/MS and database search, **Figures 3, S17**, and **S18**. We identified alpha enolase dimer and Glutathione S-transferase P dimer from the HeLa cell lysate sample, **Figure 3**. Those two protein complexes were detected with clean mass spectra with average masses of 94,440 Da and 46,440 Da, respectively. The annotated MS/MS spectra of two parent m/z ions show many matched b and y types of fragment ions from higher-energy collisional dissociation (HCD). The fragmentation patterns of the two identified proteins, alpha enolase and Glutathione S-transferase P, indicate a series of backbone cleavages near the N- and C-terminal regions, ensuring the high confidence of protein identifications. The mass of alpha enolase dimer (94,440 Da) is 364 Da heavier than two alpha enolase molecules (47038×2=94,076 Da), which may correspond to the four Mg^2+^ cofactors (96 Da) and post-translational modifications (PTMs) in the middle region of the protein sequence, i.e., phosphorylation and acetylation, according to the information in the UniProt database. The mass of Glutathione S-transferase P dimer (46,440 Da) is close to the mass of two Glutathione S-transferase P molecules (23225×2=46,450 Da).

**Figure 3.**
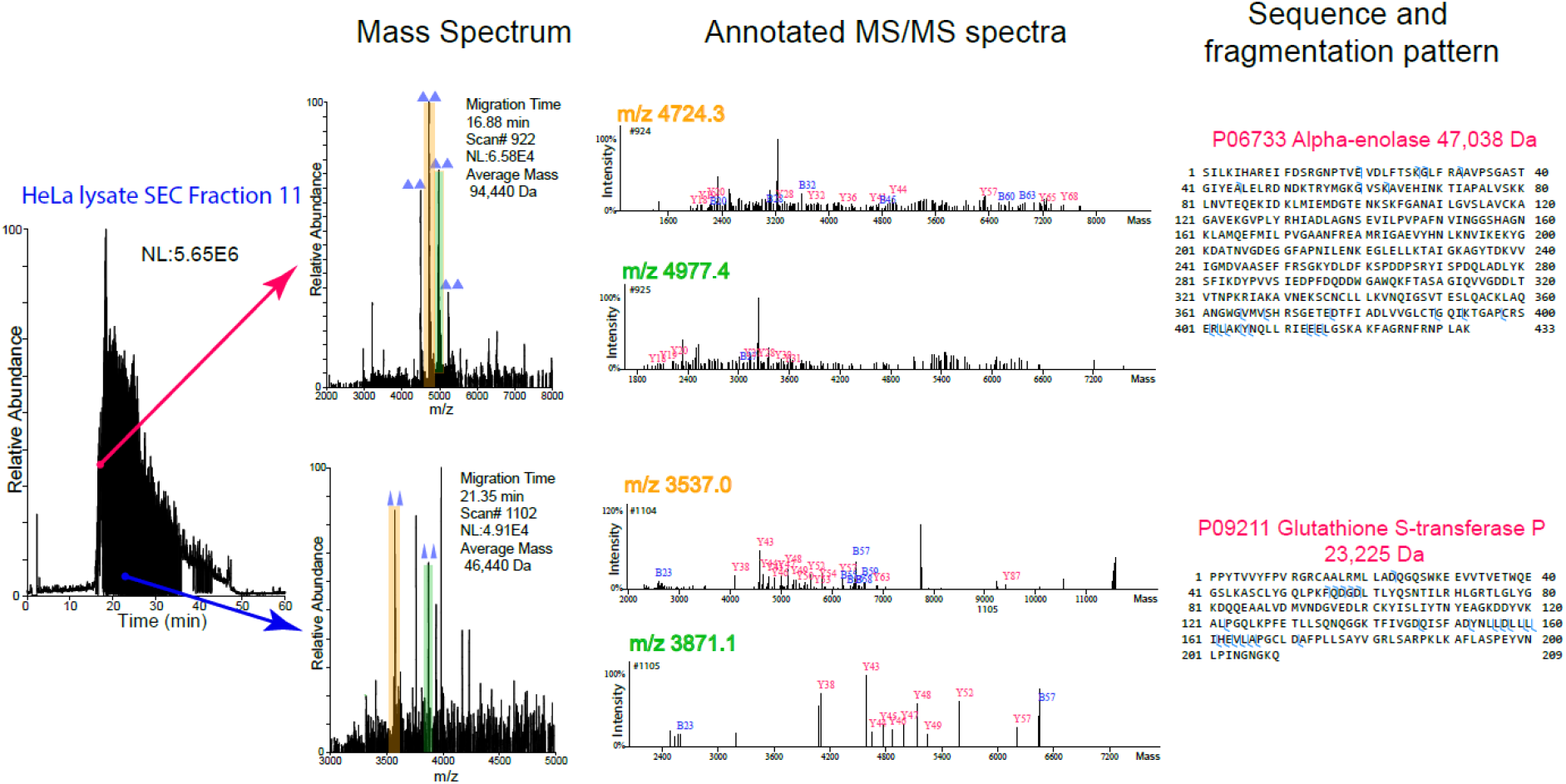
Example complexoforms (dimers) identified from CZE-MS/MS analysis of one SEC fraction of a HeLa cell lysate. The mass spectra, annotated MS/MS spectra, and fragmentation patterns of the monomers of complexoforms from higher-energy collisional dissociation (HCD) are shown. For the annotated MS/MS spectra, the data of two different charge states (two different m/z ions) of each complexoform are shown.

Similarly, CZE-MS/MS identified Glucose-6-phosphate isomerase dimer and enolase dimer in the *E. coli* cell lysate sample, **Figure S17**. The mass of the Glucose-6-phosphate isomerase dimer from MS analysis (123,090 Da) matches closely with the mass of two protein molecules (123,060 Da). The mass of enolase dimer from our measurement (91,120 Da) is 72 Da heavier than two enolase molecules (91,048 Da), which may correspond to the Mg^2+^ cofactors, according to the UniProt database. For the human plasma samples, we identified multiple complexoforms of immunoglobulin and albumin, **Figure S18**. For albumin, we detected monomer (66,440 Da), dimer (133,020 Da), trimer (199,455 Da), and tetramer (266,105 Da). The monomer mass is close to the theoretical mass of albumin (66,472 Da), only based on the protein sequence. Albumin could have many cofactors and PTMs. The mass differences between dimer and 2×monomer, trimer and 3×monomer, and tetramer and 4×monomer should be due to these cofactors and PTMs. For the immunoglobulin, using only the MS/MS spectra by database search, we identified a protein, immunoglobulin lambda variable 7-46, suggesting that the protein signal in the mass spectrum with a mass of 79550 Da (m/z 4420.4, +18) should be related to immunoglobulin. Based on our BUP data of the human plasma sample [44], there are no immunoglobulin proteins or other proteins with a mass close to 79,550 Da, only considering the most abundant proteins (top 100) in the human plasma sample. We speculate that the 79,550-Da protein could be half of an immunoglobulin protein in the human plasma, which is typically about 150-190 kDa in mass. The 159,150 Da, 246,118 Da, 326,875 Da proteins could be the intact immunoglobulin protein (79,550×2=159,100 Da), a complexoform containing the intact immunoglobulin protein and an 87-kDa protein, and a complexoform containing the dimer of the immunoglobulin protein and an 87-kDa protein, respectively. More additional studies are needed to confirm the identities of the immunoglobulin protein and the 87-kDa protein.

The pilot CZE-MS/MS data here indicate the high potential of large-scale native proteomics in discovery mode for complexoform identifications in complex biological samples using our reproducible native CZE-MS/MS technology operated in the DDA mode.

In summary, our solid bottom-up, top-down, and native proteomics data from about 170 CZE-MS runs (∼170 hours of instrument time) provide comprehensive evidence for the long-term reproducibility of CZE-MS/MS in analyzing complex mixtures of peptides, proteoforms, and complexoforms. The results indicate that the CZE-MS technology is robust enough for broad adoption in proteomics applications. For native proteomics, native CZE-MS achieved reproducible and highly sensitive measurement of complexoforms in various complex biological samples. We expect that native CZE-MS/MS operated in DDA mode will be a powerful tool for global discovery native proteomics with the assistance of advanced bioinformatics tools for complexoform identification and quantification through database search. More efforts in organizing hands-on training activities focusing on CZE-MS, such as CZE-MS summer schools, will be crucial to accelerate the broad adoption of CZE-MS technology.

## Acknowledgements

The authors thank the support from the National Cancer Institute (NCI) through grant R01CA247863 and the National Institute of General Medical Sciences (NIGMS) through grant R35GM153479.

## Conflict of Interests

Xiaowen Liu has a project contract with Bioinformatics Solutions Inc., a company that develops software for MS data processing.

## Supporting Information

The experimental details and supporting figures are included in Supporting Information I. The lists of detected complexoform masses and identified PrSMs by database search from *E. coli* cells, HeLa cells, and human plasma are included in Supporting Information II. The authors have cited additional references within the Supporting Information I.

## Author Contributions

Guijie Zhu collected all the multi-level proteomics data used in the manuscript, analyzed the data, and wrote the first draft of the manuscript. Yifan Yue helped with the data analysis. Jorge A Colón Rosado helped with the HeLa cell lysate sample preparation. Xiaowen Liu carried out the database searches of some native proteomics data. Liangliang Sun designed the experiments, oversaw the project, and edited the manuscript. All authors made comments on the manuscript.

## Data Availability Statement

The MS raw files and database search results of BUP, TDP, and native proteomics have been deposited to the ProteomeXchange Consortium via the PRIDE repository [46] with the dataset identifier PXD073607.

