## Supporting Information I for "Decades of dreams coming true: capillary zone electrophoresis-mass spectrometry for reproducible multi-level proteomics"

**Experimental procedure**

***Materials and chemicals***

Bare fused silica capillaries (50-μm i.d., 360-μm o.d.) were purchased from Polymicro Technologies (Phoenix, AZ). 3-(Trimethoxysilyl) propyl methacrylate, ammonium persulfate, ammonium acetate (NH_4_Ac), Dulbecco’s phosphate-buffered saline (dPBS), were purchased from Sigma-Aldrich (St. Louis, MO). Protease inhibitors (cOmplete ULTRA Tablets), phosphatase inhibitors (PhosSTOP) and HeLa protein digest standard, hydrofluoric acid (HF), LC/MS grade water, methanol and MilliporeSigma™ Amicon™ Ultra-0.5 Centrifugal Filter 10 kDa units for buffer exchange were purchased from Fisher Scientific (Pittsburgh, PA). Acrylamide were purchased from Acros Organics (NJ, USA).

***Sample preparation***

**Cell Culture**

*E. coli* (K12_MG1655 strain) was cultured in Terrific Broth (TB) medium at 37 °C until OD600 reached 0.7. The *E. coli* cells were washed with PBS three times.

HeLa cells were cultured in MEM-containing 10% fetal bovine serum (v/v) and maintained in a humidified atmosphere of 95% air and 5% CO_2_ at 37 °C. The adherent cell layer was washed with PBS and then trypsinized with 0.05% trypsin-EDTA solution for 5 min at 37 °C. Then, the cells were centrifuged at 250 × g for 5 min to remove trypsin, followed by washing with PBS three times.

**BUP sample preparation**

Commercialized HeLa protein digest standard (20 µg) was first dissolved in 50 µL LC-MS grade water. Then, 10 µL HeLa protein digests were mixed with 10 µL 100 mM NH_4_Ac for CE-MS sample injection.

**TDP sample preparation**

0.35 g of wet *E. coli* cell pellet was suspended in 2 mL of 8 M urea containing protease and phosphatase inhibitors (5 mg/mL each). The suspended cells were then sonicated for 2 min twice in an ice bath with a Branson Sonifier 250 (VWR Scientific, IL, USA). After that, the cell lysate was centrifuged with an Allegra X-30R Centrifuge (Beckman Coulter, CA, USA) at 14000 g for 25 min. The concentration of the collected supernatant was determined using a bicinchoninic acid (BCA) assay (Thermo Fisher Scientific, MA, USA). The *E. coli* cell lysate was buffer-exchanged to 100 mM ammonium bicarbonate using 10 kDa cut-off Amicon Ultra centrifugal filters. Finally, the *E. coli* sample was diluted to 1.5 mg/mL using 100 mM ammonium bicarbonate for CZE-MS analysis.

HeLa cell proteins were extracted using 8 M urea containing protease and phosphatase inhibitors (5 mg/mL each). 5 × 10^6^ HeLa cells were suspended in 300 µL lysis buffer and sonicated on ice bath for 2 min with a Branson Sonifier 250 at 15 s on/off intervals for 10 times. The lysate was then centrifuged with an Allegra X-30R Centrifuge (Beckman Coulter, CA, USA) at 16,000 × g for 15 min at 4 °C, and the supernatant containing the extracted proteins was collected. Protein concentration was determined to be 6 mg/mL using a BCA assay. Then, 36 µL of 500 mM dithiothreitol (DTT) was added to reduce proteins at 35 °C for 1 h. 1 mg of HeLa cell proteins was first loaded on 100-kDa molecular weight cutoff (MWCO) filter units. The flow-through proteins were further buffer-exchanged into 100 mM ammonium acetate (NH_4_Ac, pH=7.5) using 10-kDa molecular weight cutoff (MWCO) filter units. Totally, 70 µL HeLa cell proteins (3 mg/mL) were collected for CZE-MS/MS.

**Native proteomics sample preparation**

**E.coli cell lysate preparation**

About 200 mg *E. coli* cell pellet (wet weight) was suspended in 2 mL of Dulbecco’s phosphate-buffered saline (DPBS, Sigma-Aldrich, MO, USA) buffer, and then sonicated in an ice bath with a Branson Sonifier 250 for 5 minutes. After that, the lysates were centrifuged at 16,000 g for 15 minutes to collect the supernatant. Then, the protein concentration was determined as 4 mg/mL by using a BCA assay. The samples were aliquoted to 100 µL of solution (400 µg protein) each and stored at -20 °C for use.

180 µL of the protein solution (~ 700 µg proteins) was buffer-exchanged using 10 kDa molecular weight cutoff (MWCO) filter units. Briefly, 200 µL of 100 mM NH_4_Ac was used to condition the filter first. Then the protein solution was buffer-exchanged to 100 mM NH_4_AC by washing with 100 mM NH_4_AC (150 µL) for 4 times and then was buffer- exchanged to 20 mM NH_4_Ac by washing with 20 mM NH_4_Ac for 3 times. Finally, 70 µL of 20 mM NH_4_Ac was added to the filter. About 100 µL protein solution was collected by inverting the filter unit and centrifuge. The protein solution was used for native CZE-MS analysis.

**Native HeLa cell lysate preparation**

HeLa cell proteins were extracted using DPBS. 3 mL of DPBS containing protease and phosphatase inhibitors (15 mg each) was used. The sample was divided equally into two 1.5 mL vials to generate technical duplicates. After the cell pellet was resuspended in DPBS, the resulting duplicates were lysed on ice for 20 min by sonication with a Branson Sonifier 250 (VWR Scientific, IL, USA) at 15 s on/off intervals. The lysate was then centrifuged in an Allegra X-30R Centrifuge at 14,000 × g for 30 min at 4 °C, and the supernatant containing the extracted proteins was collected. Duplicates were combined, and protein concentration was determined to be 3.3 mg/mL using a BCA assay.

The DPBS lysate was buffer-exchanged into 100 mM NH_4_Ac (pH=7) using 30 kDa molecular weight cutoff (MWCO) filter units. 1 mL of HeLa lysate was loaded onto three different filter units and centrifuged at 14,000 × g for ~30 min at 4 °C until concentrated to less than 50 µL. Subsequently, 200 µL of 100 mM NH_4_Ac was added, mixed by gentle pipetting, and centrifuged under the same conditions. This wash step was repeated three times, and the final retentate was collected with a micropipette. The three replicates were combined, and protein concentration was determined to be 2 mg/mL with a BCA assay. The sample was fractionated by size exclusion chromatography (SEC).

SEC fractionation was performed using a 1260 Infinity II HPLC (Agilent, CA, USA) system equipped with a UV–Vis detector (250 nm). 100 mM NH_4_Ac was used as the mobile phase for separation. 20 μL of sample was injected into an Agilent column (4.6 × 300 mm, 5 μm, 500 Å) at a flow rate of 200 μL/min. Fractions were collected every 1 min over a 40 min separation, beginning at the first UV-Vis absorbance peak. Seven injections (replicates) were performed, and the fractions collected at the same elution time were combined. Each combined fraction was concentrated using a 30-kDa MWCO filter unit by centrifugation at 14,000 × g for approximately 30 min at 4 °C, until the final volume was reduced to ~30 µL. The concentrated HeLa protein fractions were stored at -20 °C until further analysis. In this specific experiment, only one SEC fraction (fraction 11) was used.

**Native human plasma preparation**

Commercial healthy human plasma (Innovative Research, MI, USA) was first diluted by 100 mM NH_4_Ac by a factor of 10, then 200 uL of diluted human plasma was buffer- exchanged to 20 mM NH_4_Ac by using 30 kDa MWCO filter units. After that, the final volume was changed to 100 µL by adding a corresponding volume of 20 mM NH_4_Ac.

***Preparation of LPA-coated separation capillary***

The inner wall of the separation capillary (50-μm i.d., 360-μm o.d.) was coated with linear polyacrylamide (LPA) based on the protocol described in previous references. [1,2] Briefly, a bare fused silica capillary was successively flushed with 1 M hydrochloric acid, water, 1 M sodium hydroxide, water, and methanol, followed by treatment with 3-(trimethoxysilyl) propyl methacrylate for at least 48 hours to introduce carbon-carbon double bonds on the inner wall of the capillary. The treated capillary was filled with degassed acrylamide solution in water (4%) containing 0.035% ammonium persulfate, followed by incubation in a 50 °C water bath for 60 min with both ends sealed by silica rubber. After that, the capillary was flushed with water to remove the unreacted reagents. Then, one end of the LPA-coated capillary was etched with HF based on the protocol in reference [3] for 70 minutes to reduce its outer diameter to around 100 μm.

***CZE-MS***

**CZE-MS of BUP and TDP**

For both BUP and TDP, a Beckman CESI8000 Plus CE autosampler(Beckman Coulter) was used for the automated operation of CZE. A home-built electrokinetically pumped sheath flow interface was used to couple CZE to an Orbitrap Exploris 480 mass spectrometer. [4,5] A 95-cm LPA-coated capillary (50 μm i.d., 360 μm o.d.) was used for BUP analysis, and a 100-cm LPA-coated capillary (50 μm i.d., 360 μm o.d.) was used for TDP analysis. For CZE-MS, nanoelectrospray ionization (nanoESI) emitters with an orifice size of 30~35 μm were pulled from borosilicate glass capillaries (1.0 mm o.d., 0.75 mm i.d.) using a Sutter P-1000 flaming/brown pipette puller (Sutter Instrument, Novato, CA) for ESI spray, and the voltage of nanoESI was set to +2 kV. The BGE consisted of 5% acetic acid (v/v), while the sheath liquid contained 0.2% formic acid and 10% methanol (v/v). 60 nL of each sample was injected for analysis using 5 psi air pressure calculated based on Poiseuille’s law. For CZE separation, a high voltage of +30 kV was applied for 60 minutes (BUP), 70 minutes (TDP), followed by 10 minutes (BUP) and 15 minutes (TDP) of flushing method with +30 kV and 20 psi forward.

For BUP, experiments were performed in a data-dependent acquisition (DDA) mode. Full MS scans were acquired in the Orbitrap mass analyzer over the m/z 300–1,500 range with a resolution of 60,000 (at 400 m/z). The 15 most intense peaks with a charge state of 2-6 were fragmented in the higher-energy collisional dissociation (HCD) cell with collision energy of 28% and analyzed by the Orbitrap mass analyzer with a resolution of 30,000 (at 200 m/z). The intensity threshold was 5.0e4, the isolation window was 2 m/z, and the dynamic exclusion duration was 15 s. Normalized AGC target on MS1 was 300%, normalized AGC target on MS2 was 100%, and a single microscan for both MS1 and MS2.

For TDP, the full MS parameters included a high mass resolution of 480,000 (at m/z 200) with a single microscan, covering a scan range of 600-2000 m/z. Precursor ions were isolated with a 2 m/z window and subjected to fragmentation through higher-energy collisional dissociation (HCD) with a normalized collision energy of 25%. Only precursor ions with an intensity exceeding 5E4 and a charge state ranging from 3 to 60 were selected for fragmentation. Product ions were detected with a resolution of 60,000 (at 200 m/z), utilizing a single microscan, and maintaining a normalized AGC target value of 300% for both conditions. Dynamic exclusion was enabled with a duration of 15 seconds and a mass tolerance of 10 ppm (parts per million). Additionally, the "Exclude isotopes" function was activated.

**CZE-MS of native proteomics**

CZE-MS experiments were performed by coupling a CMP EXE-001 CE autosampler (CMP Scientific, Brooklyn, NY) to a Q-Exactive Ultra-High Mass Range (UHMR) Orbitrap mass spectrometer (Thermo Fisher Scientific, Waltham, MA) through a commercial electrokinetically pumped sheath flow nanospray interface (EMASS-II, CMP Scientific, Brooklyn, NY). [4,5] The nanoelectrospray ionization (nanoESI) emitters used in the EMASS-II interface were pulled from borosilicate glass capillaries (1.0 mm o.d., 0.75 mm i.d.) using a Sutter P-1000 flaming/brown pipette to an orifice size of 30~35 μm. The voltage of nanoESI was set to +2 kV. An LPA-coated capillary (50 μm i.d., 360 μm o.d.) was employed with one end etched to around 100 µm by HF (90 cm for *E.coli* and HeLa, 85 cm for Human Plasma). Both background electrolyte (BGE) and the sheath liquid contained 20 mM NH_4_Ac (pH 7.5). Sample injection into the capillary was performed using 200 mbar air pressure, with 70 nL injection calculated based on Poiseuille’s law.

For *E. coli* high concentration sample (~3 mg/mL), a high voltage of +26 kV with an assisting 60 mbar was applied for 5000 seconds, followed by 1500 mbar applied for 1000 seconds for capillary flushing. For *E. coli* low concentration sample (~1 mg/mL), a high voltage of +20 kV with an assisting 80 mbar was applied for 40 minutes, followed by 800 mbar applied for 15 minutes for capillary flushing. For the HeLa cell lysate SEC fraction, a high voltage of +20 kV with an assisting 80 mbar was applied for 60 minutes, followed by 1000 mbar applied for 20 minutes for capillary flushing. For human plasma sample, a high voltage of +25 kV with an assisting 1 psi was applied for 70 minutes, followed by 20 psi applied for 20 minutes for capillary flushing.

For the MS settings, transfer capillary temperature was 300 °C. The full MS parameters included the mass range for MS scans 2000-15000 m/z, the resolution 6,250 defined at 400 m/z, AGC target: 1E6, maximum injection time: 200 ms, in-source trapping (IST) desolvation voltage -100 V, trapping gas setting 7, and 5 microscans. For the experiment applied MS2, a DDA mode was applied. The six most intense ions were selected sequentially for MS/MS. Precursor ions were isolated with a 50 m/z window and subjected to fragmentation through higher-energy collisional dissociation (HCD). A normalized HCD collision energy ranging from 30% to 300% was tested to maximize the complexoform fragmentation. Only precursor ions with an intensity exceeding 5E4 and a charge state higher than 5 were selected for fragmentation. Product ions were detected in either a low mass resolution or a high mass resolution. For the low-mass-resolution condition, a mass resolution of 6,250 (at 400 m/z) was used with 10 microscans, and a normalized AGC target value of 1E6. For the high- mass-resolution condition, a mass resolution of 100,000 (at m/z 400) was used.

***Data Analysis***

For the BUP experiment, the raw files were processed using MaxQuant software(Version 2.6.7.0).[6] The MS/MS spectra were searched against the UniProt human proteome database (uniprotkb_proteome_UP000005640_2025_04_07_HUMAN). The peptide mass tolerances for the initial and main searches were set to 20 ppm and 4.5 ppm, respectively, with a fragment ion mass tolerance of 20 ppm. Trypsin/P was specified as the protease, carbamidomethylation of cysteine residues (+57.0215 Da) was set as a fixed modification, and dynamic modifications included oxidation on methionine and acetylation at the protein N-terminus. False discovery rates (FDRs) were controlled at 1% for both peptides and proteins.

All the TDP mass spectral proteoform identification was performed using TopPIC (Top-down mass spectrometry-based Proteoform Identification and Characterization) pipeline. [7] Briefly, the raw file was converted to centroided mzML files by MSConvert [8] using peak-picking. Then, the spectral deconvolution was performed using Top-down mass spectrometry Feature Detection [9] (TopFD) with default parameter settings. The database search was performed using TopPIC (version 1.7.4) against the UniProt proteome database of *E. coli* (uniprotkb_proteome_UP000000625_2025_04_07_ECOLI), Human (uniprotkb_proteome_UP000005640_2025_04_07_HUMAN). The parameters were set as follows: mass error tolerance of matching masses of 10 ppm, maximum number of mass shifts of unexpected modifications of 2, ranging from -500 Da to 500 Da. The FDRs were estimated using the target-decoy approach. The spectrum- level FDR cutoff was 1%, and the proteoform-level FDR cutoff was 1%.

The average mass of protein complexes in native proteomics was obtained by mass deconvolution using UniDec [10] or ESIprot [11].

Native mass spectra of protein complexes were analyzed using the TopPIC Suite [7,12] to identify candidate protein-complex identifications, followed by manual validation to keep only high-confidence identifications. Raw data files were converted to the centroided mzML files using MSConvert [8] with peak picking. Only the MS/MS spectra were deconvoluted using TopFD (version 1.8.0) [9], and MS1 spectra were excluded from data processing. The maximum charge state was set to 3 and the maximum mass was set to 20 kDa for fragment ions. All other TopFD parameters were set to default ones. The resulting deconvoluted MS/MS spectra were then searched using a modified version of TopPIC (version 1.8.0) against the corresponding protein sequence database (*E. coli*: the UniProt E. coli proteome database with 4,402 proteins, proteome ID: UP000000625, version April 07, 2025; HeLa: a human protein database with 124 high-abundance proteins identified from BUP analysis of the same SEC fraction; plasma: a human protein database with 180 high-abundance plasma proteins identified from the BUP dataset of the same sample). Because precursor masses of protein complexes were not available, only fragment masses were used for proteoform identification in TopPIC. In database searching, each MS/MS spectrum was assigned an arbitrary precursor mass of 20 kDa, and matches were accepted when the theoretical molecular weight of a candidate protein fell within the range of 0 kDa and 1020 kDa. Search parameters were set as follows: a mass error tolerance of 15 ppm for fragment matching; a maximum of one unexpected mass shift in a proteoform; an allowable mass-shift range of -1000 kDa to +20 kDa; only complete proteoforms were searched; and an E-value cutoff of 50 was used. In addition, +/-1 Da errors were allowed for matching fragments with a mass ≥ 1000 Da. Default settings were used for all other TopPIC parameters. The E-value cutoff of 50 was chosen to search for potential proteoform candidates, while manual validation was used to report only high-confidence proteoform identifications.

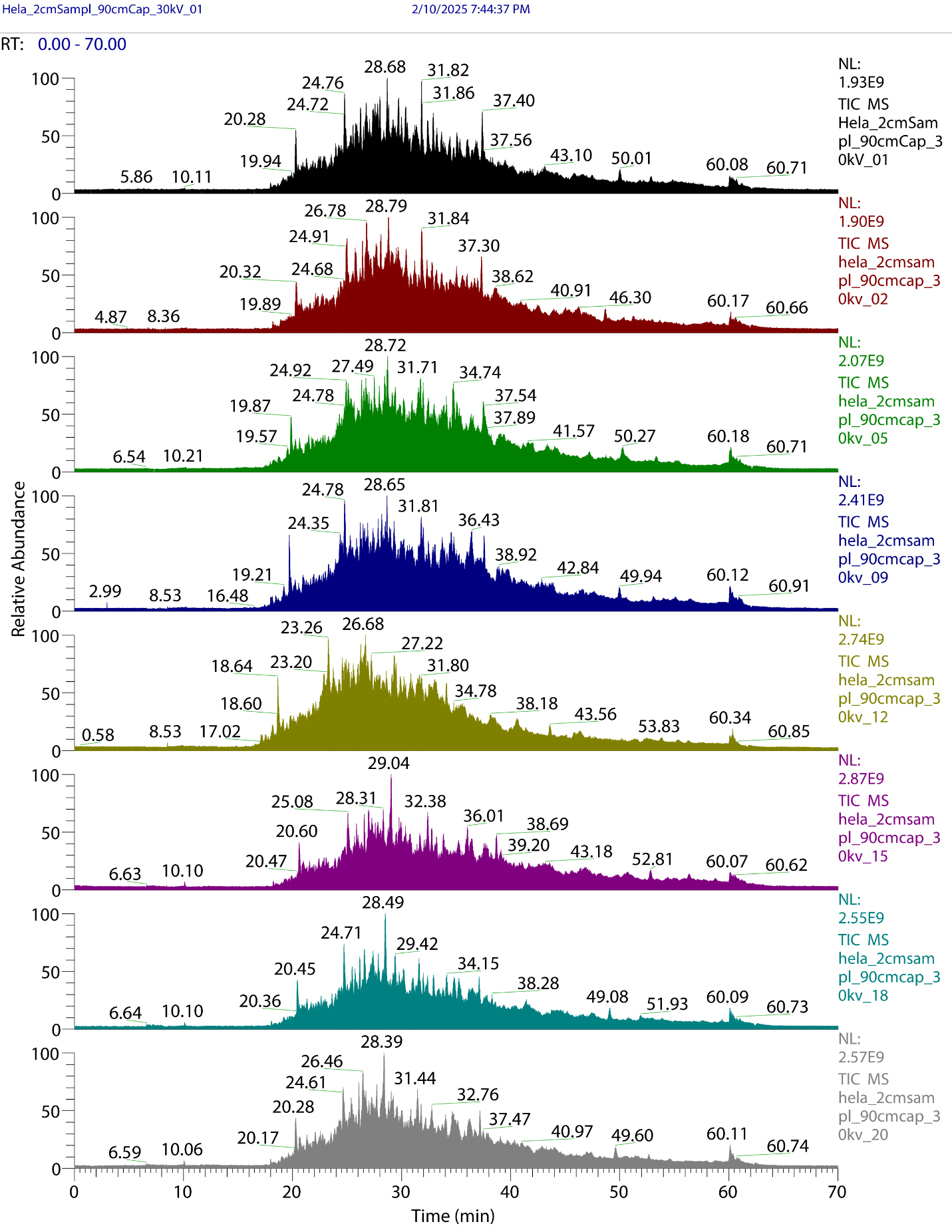

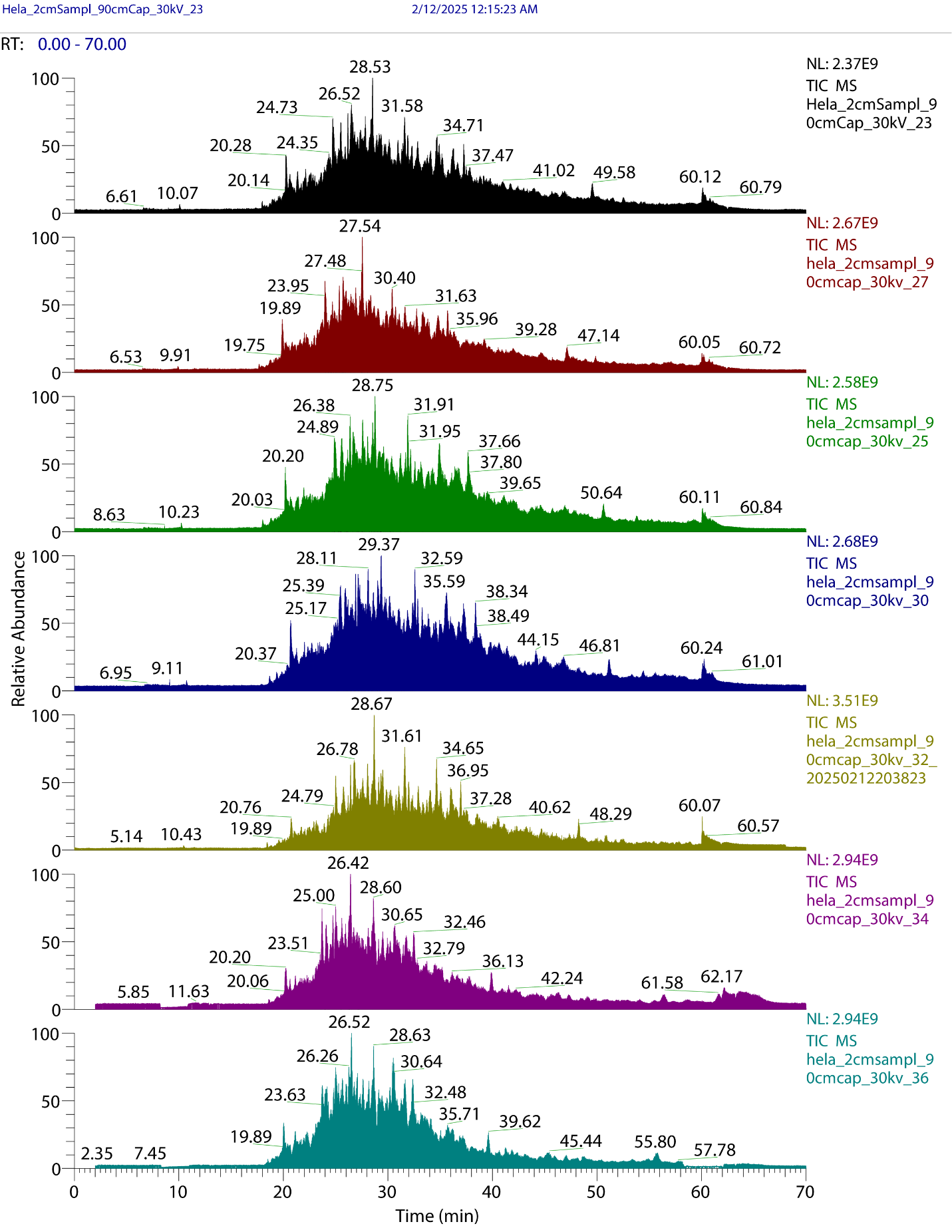

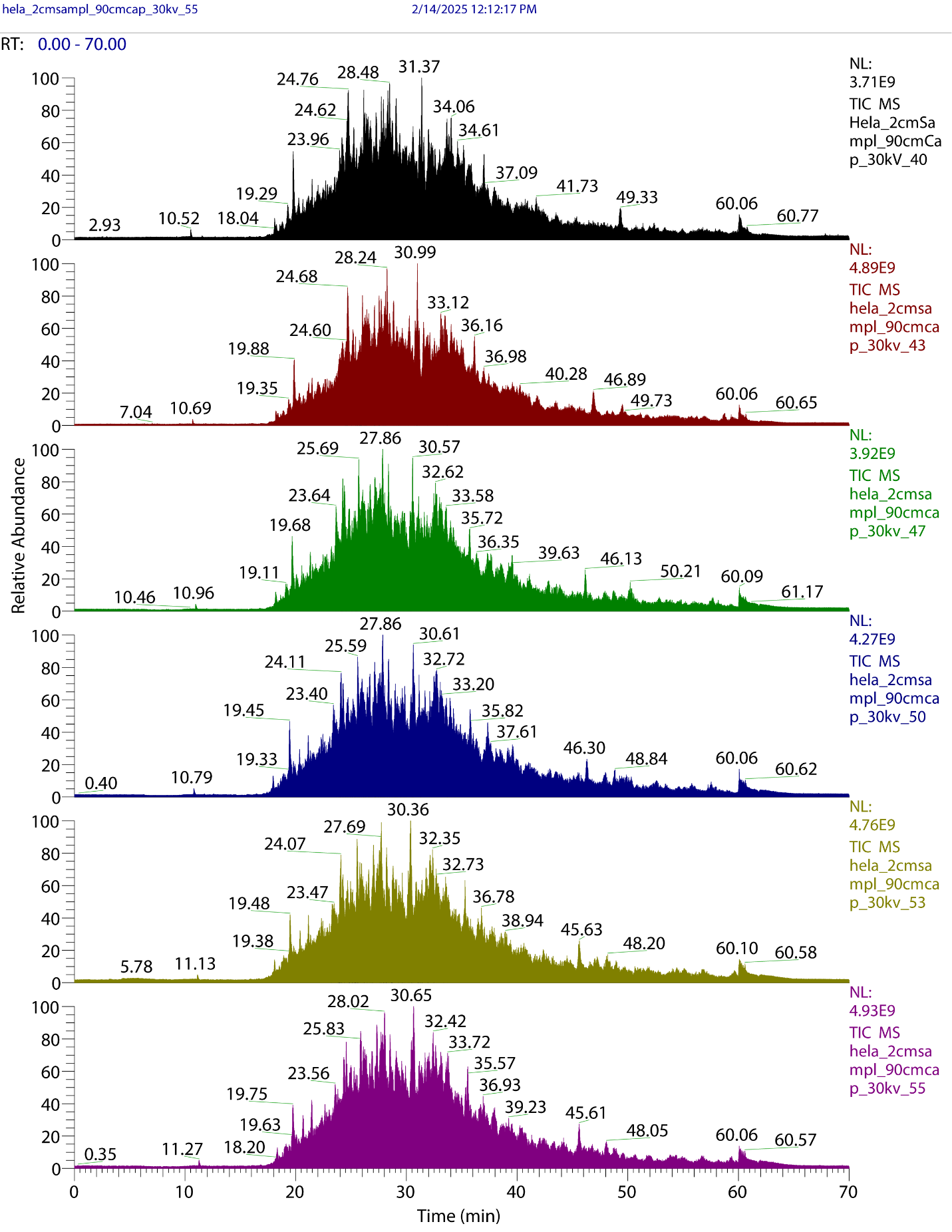

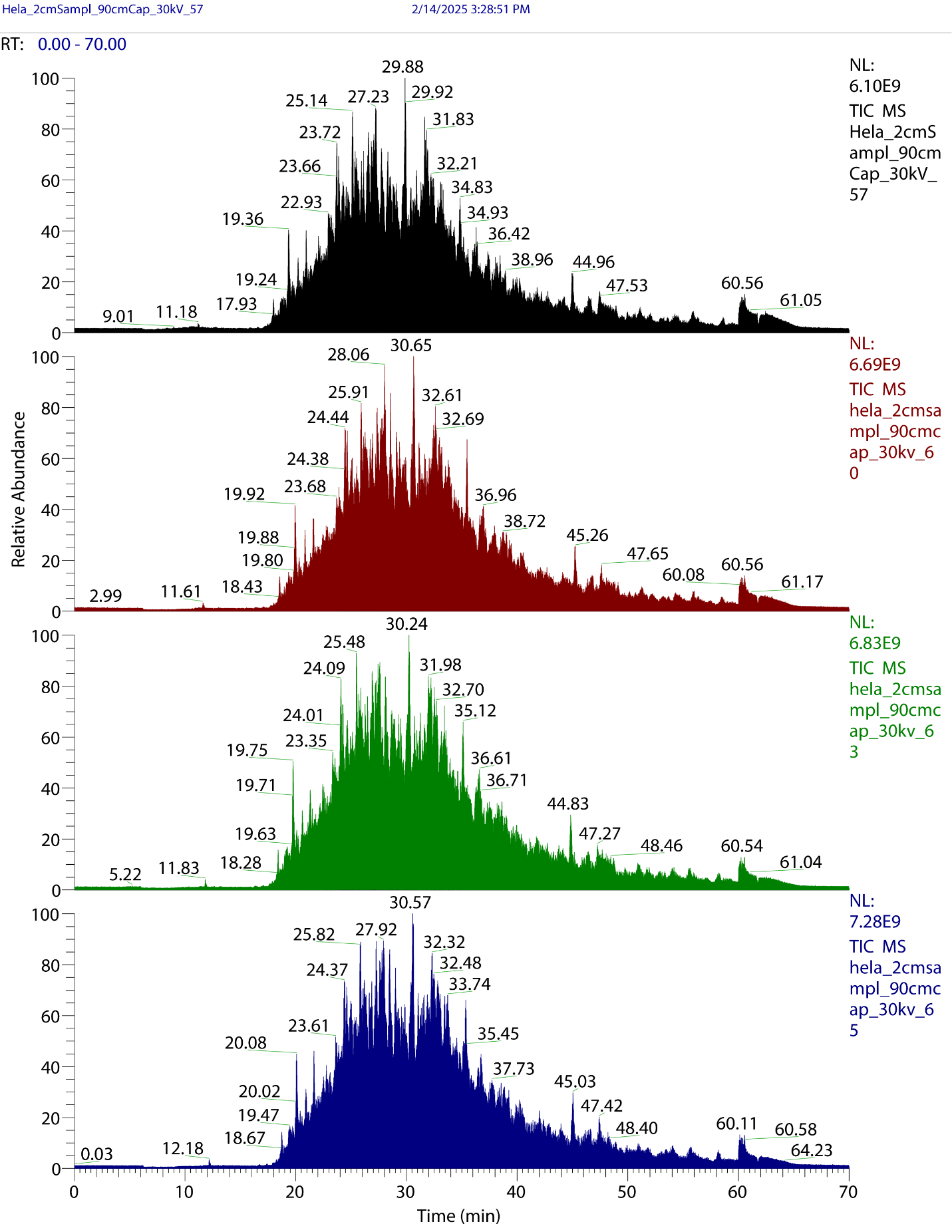

**Figure S1**. Total ion current (TIC) electropherograms of randomly selected 25 CZE-MS/MS runs (67 runs in total) of a HeLa cell digest.

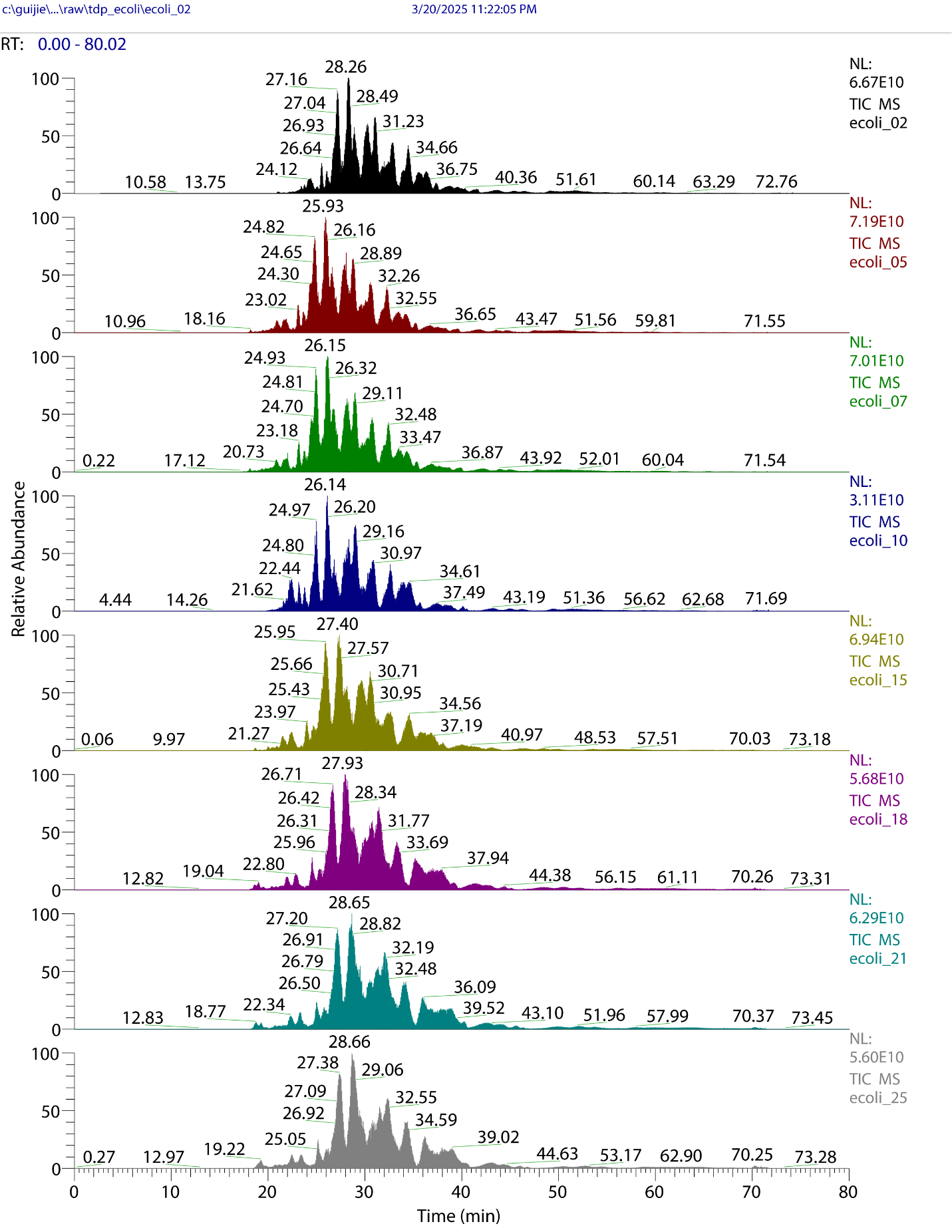

**Figure S2**. Total ion current (TIC) electropherograms of randomly selected 8 CZE-MS/MS runs (26 runs in total) of an *E. coli* cell lysate.

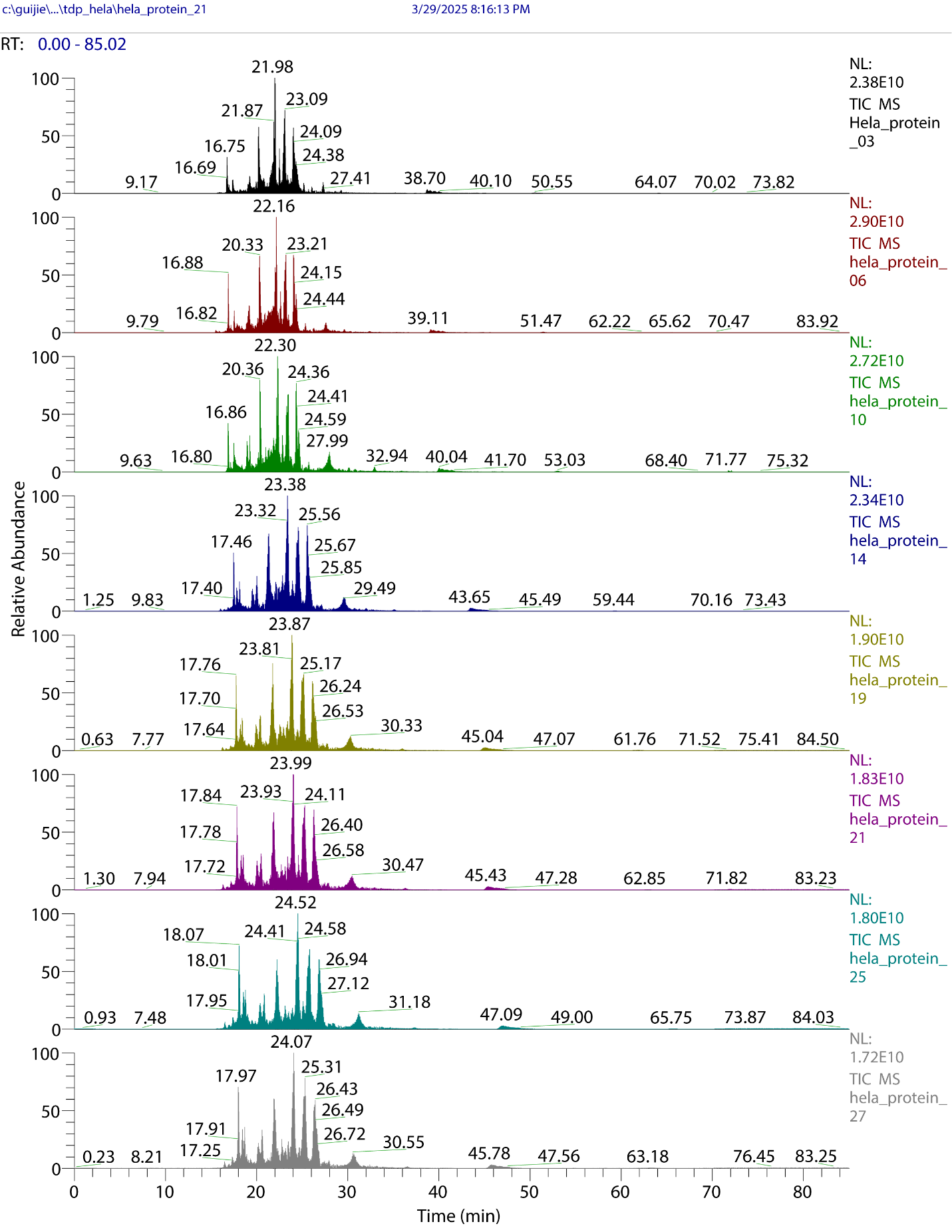

**Figure S3**. Total ion current (TIC) electropherograms of randomly selected 8 CZE-MS/MS runs (29 runs in total) of a HeLa cell lysate.

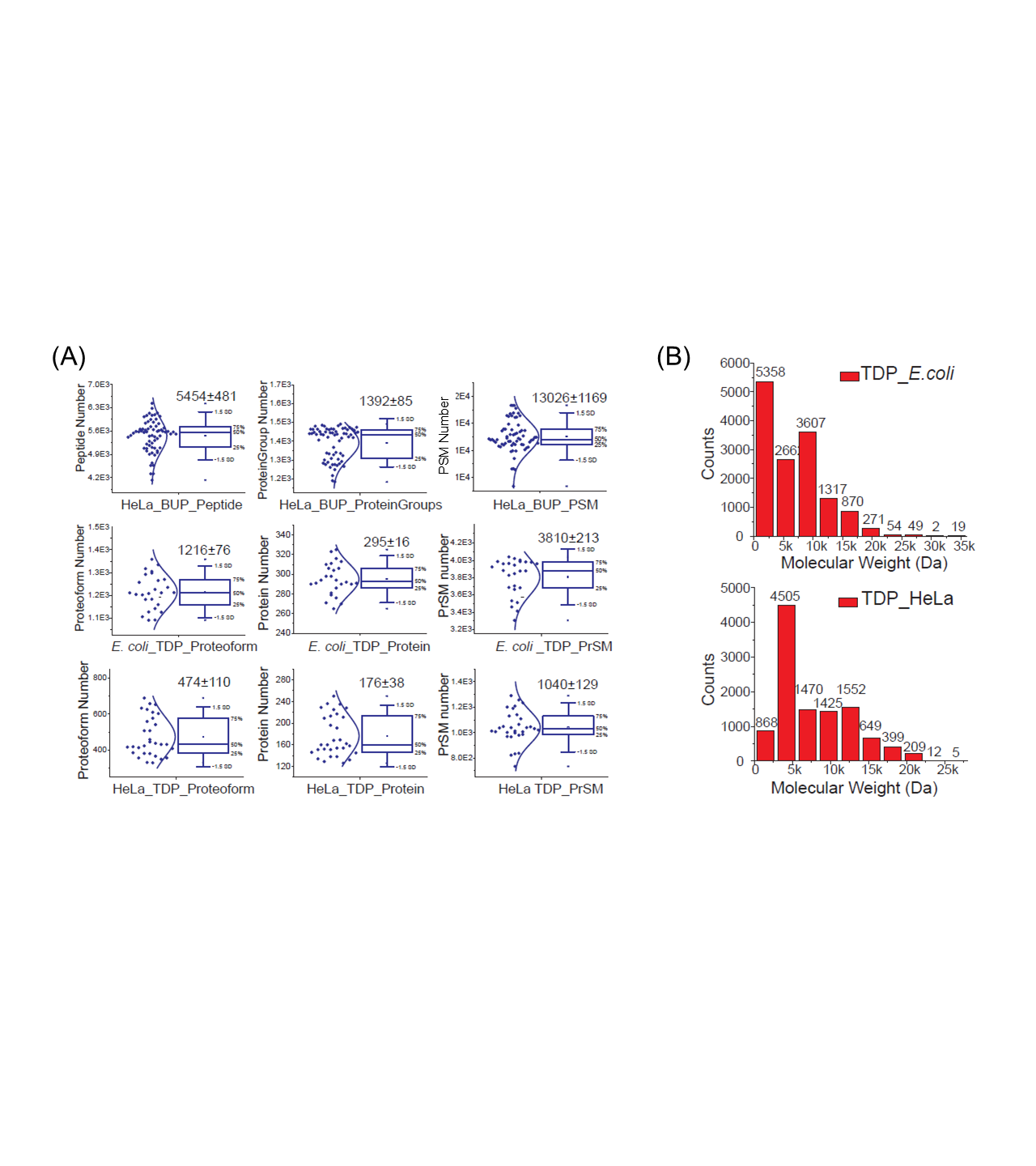

**Figure S4**. (A) Box plots of identifications of peptide spectrum matches (PSMs), peptides, and protein groups from BUP of a HeLa cell lysate, and proteoform-spectrum matches (PrSMs), proteoforms, and proteins from TDP of an *E. coli* cell lysate and a HeLa cell lysate. The numbers labelled on each figure are the mean of the number of identifications from all the CZE-MS/MS runs ± standard deviations. (B) The mass distribution of identified proteoforms from the *E. coli* sample (top) and HeLa cell sample (bottom). The combined list of proteoforms from all the CZE-MS/MS runs was used here for the figure.

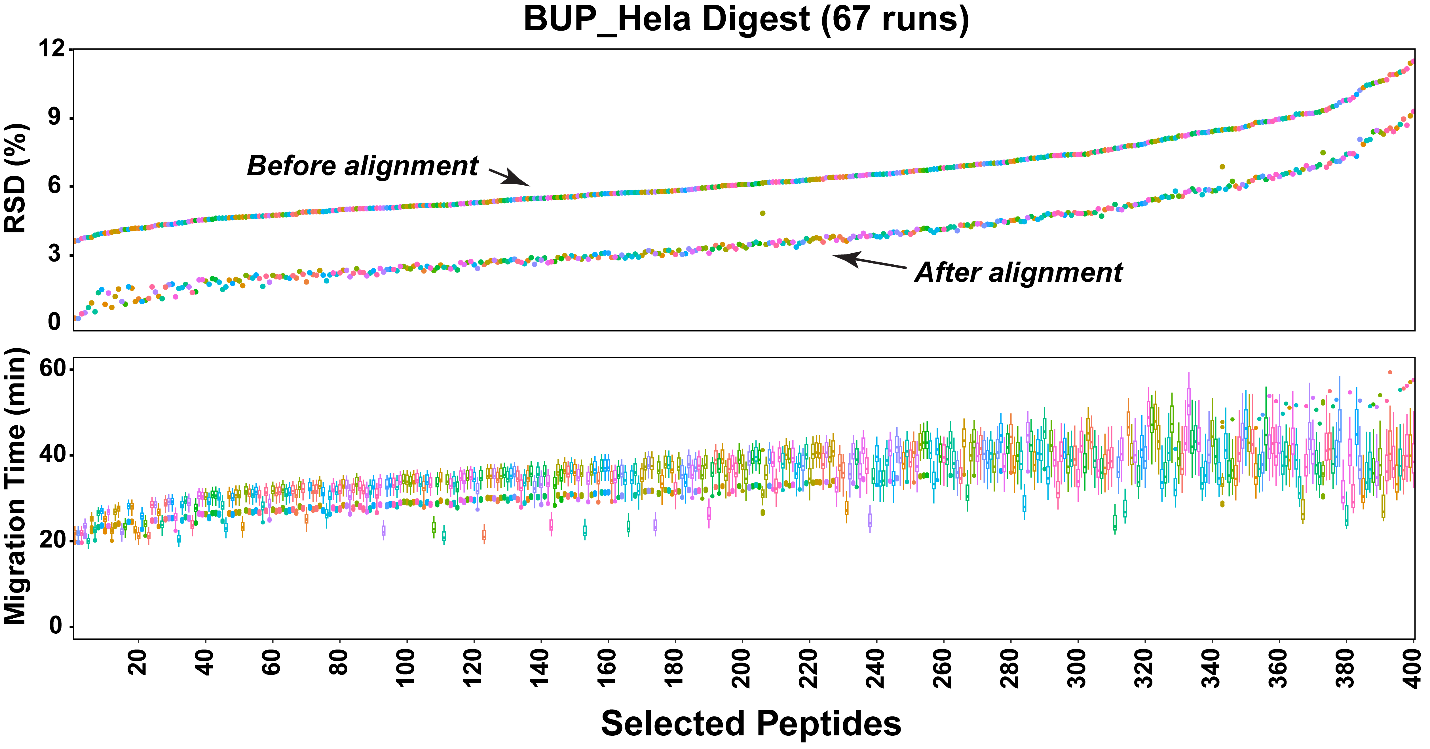

**Figure S5**. Boxplots (bottom) and relative standard deviations (RSDs) (top) of the migration time of 400 selected peptides across 67 CZE-MS/MS runs. The RSDs of peptides before and after migration time alignment are shown.

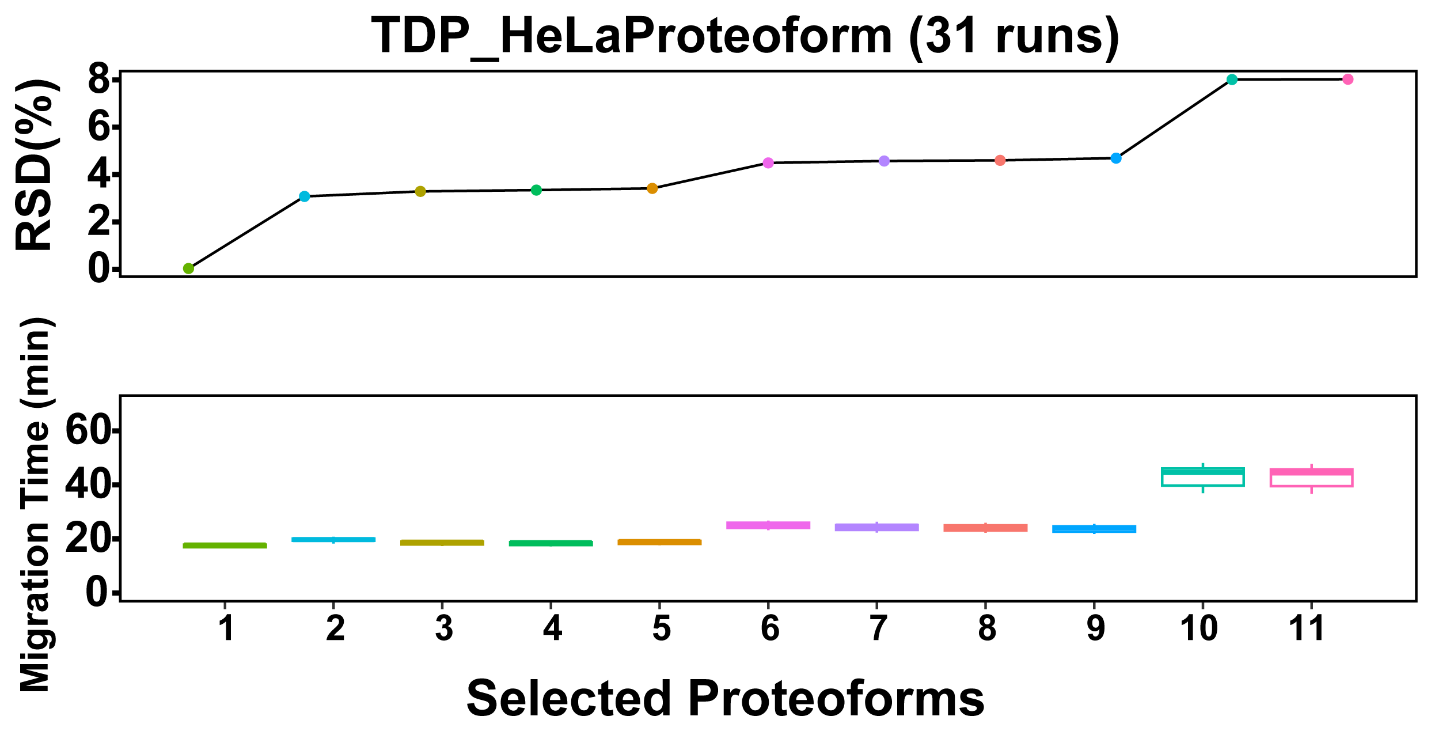

**Figure S6**. Box-plots and relative standard deviations (RSDs) of migration time of selected 11 proteoforms across 31 CZE-MS/MS runs of a HeLa cell lysate.

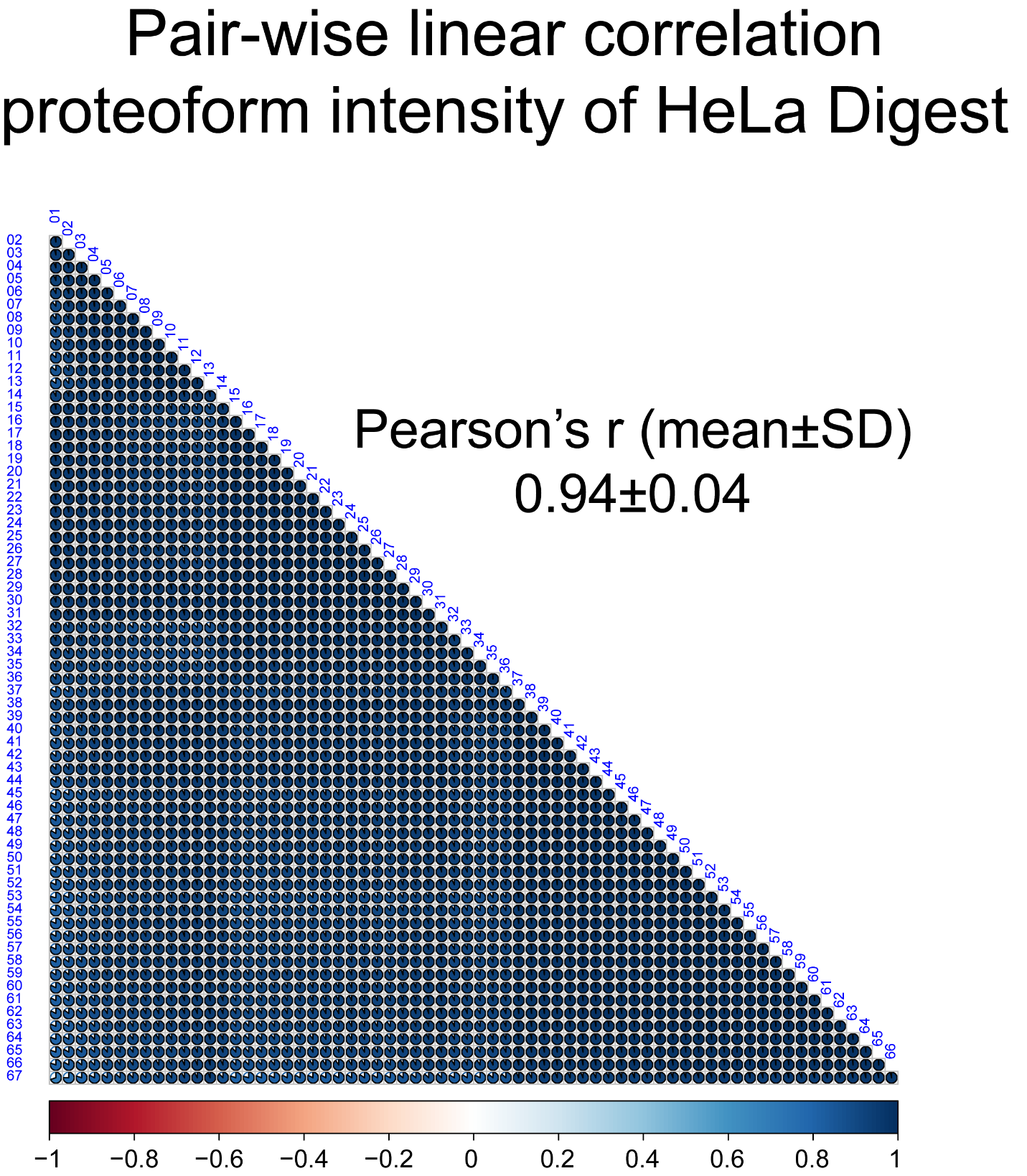

**Figure S7**. Pair-wise linear correlations of peptide intensity from CZE-MS/MS analyses of a HeLa cell digest. The color bar shows the range of Pearson’s *r* of the linear correlations. SD: standard deviation.

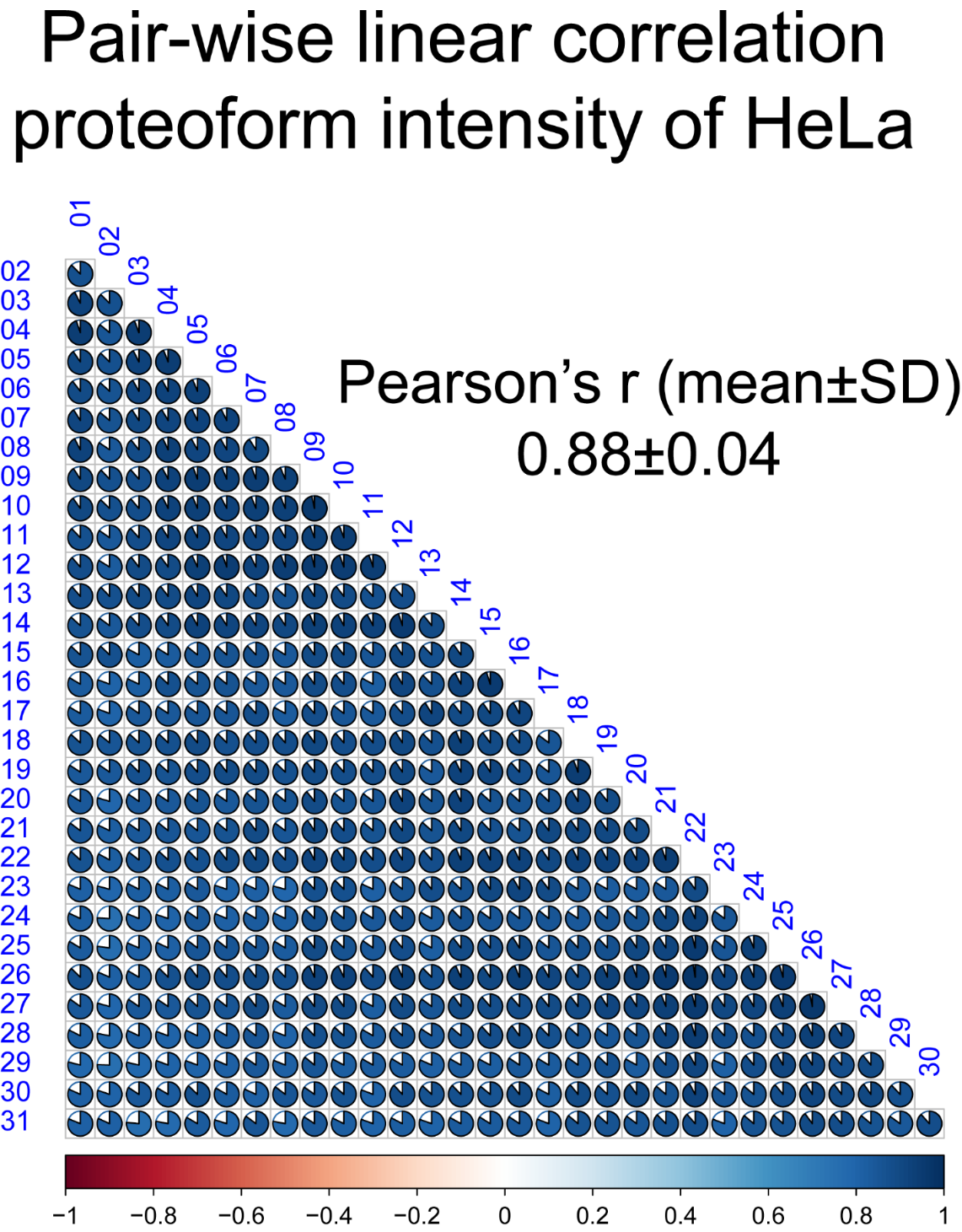

**Figure S8**. Pair-wise linear correlations of proteoform intensity from CZE-MS/MS analyses of a HeLa cell lysate. The color bar shows the range of Pearson’s *r* of the linear correlations. SD: standard deviation.

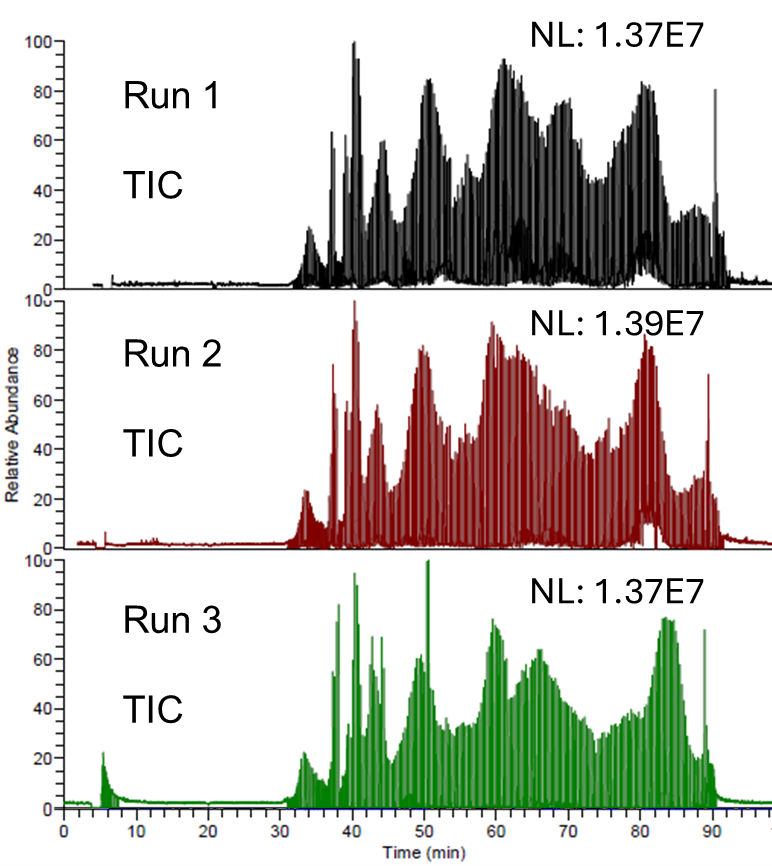

**Figure S9**. Total ion current (TIC) electropherograms of an *E. coli* cell lysate after native CZE-MS/MS analysis in technical triplicate. The MS1 spectra were acquired under a low-mass-resolution condition (6250, m/z 400), and MS/MS spectra were acquired under a low-mass-resolution condition (6250, m/z 400). The protein concentration is ~3 mg/mL.

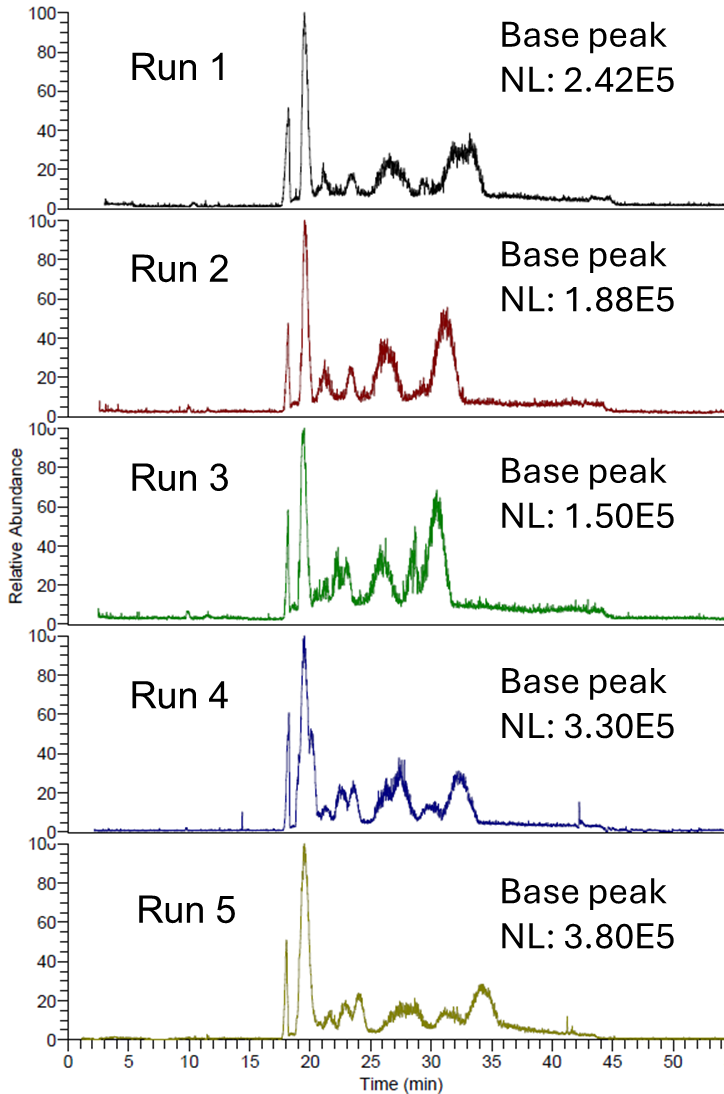

**Figure S10**. Base peak electropherograms of five native CZE-MS runs of an *E. coli* cell lysate. The protein concentration is ~1 mg/mL. Only MS1 spectra were acquired.

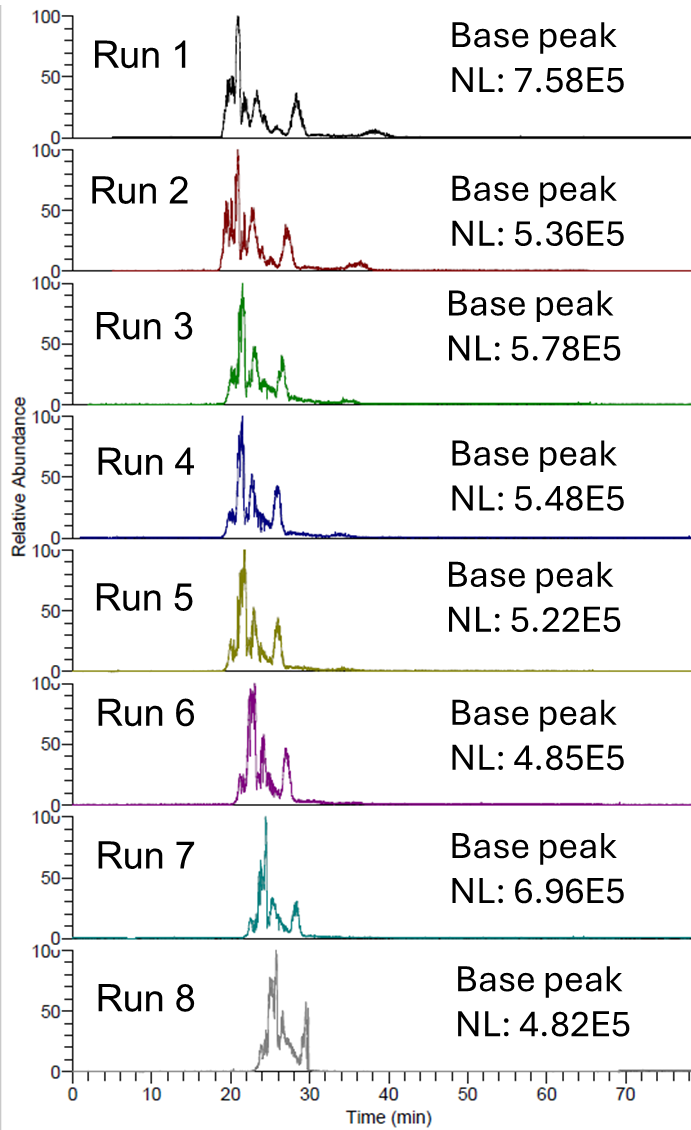

**Figure S11**. Base peak electropherograms of eight native CZE-MS runs of one size exclusion chromatography (SEC) fraction of a HeLa cell lysate. Only MS1 spectra were acquired.

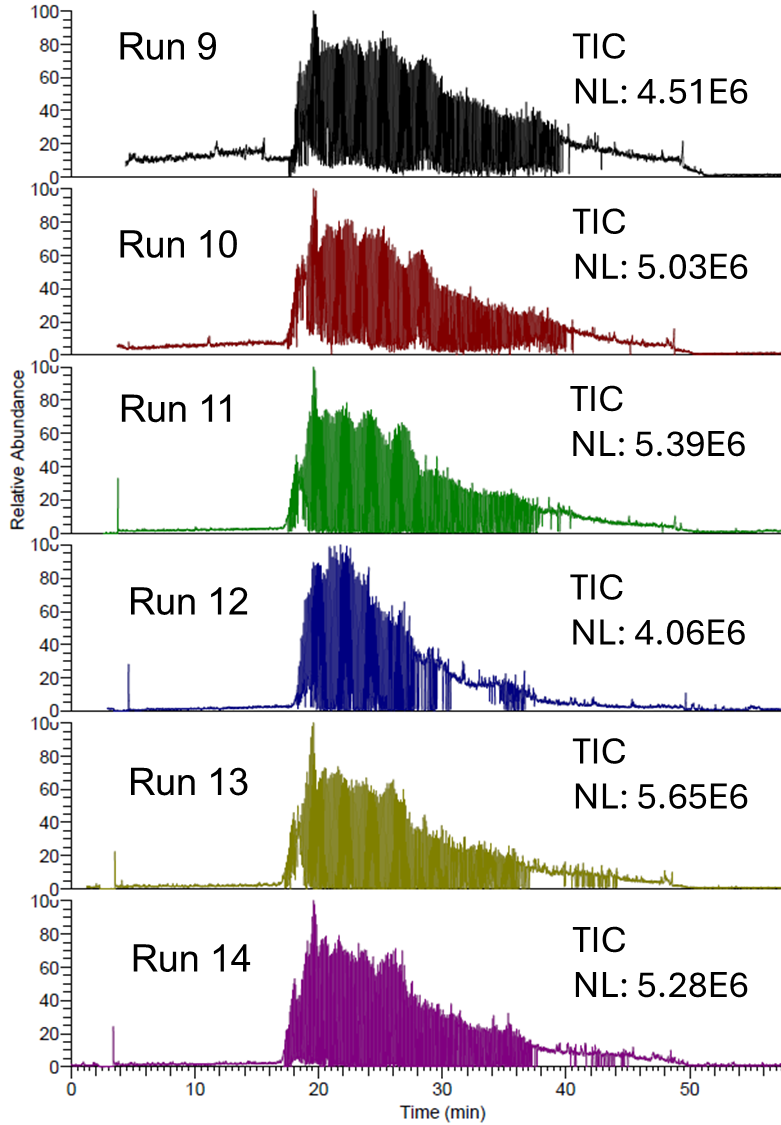

**Figure S12**. Total ion current (TIC) electropherograms of one size exclusion chromatography (SEC) fraction of the HeLa cell lysate after six native CZE-MS/MS runs. The MS1 spectra were acquired under a low-mass-resolution condition (6250, m/z 400), and MS/MS spectra were acquired under a high-mass-resolution condition (100,000 at m/z 400).

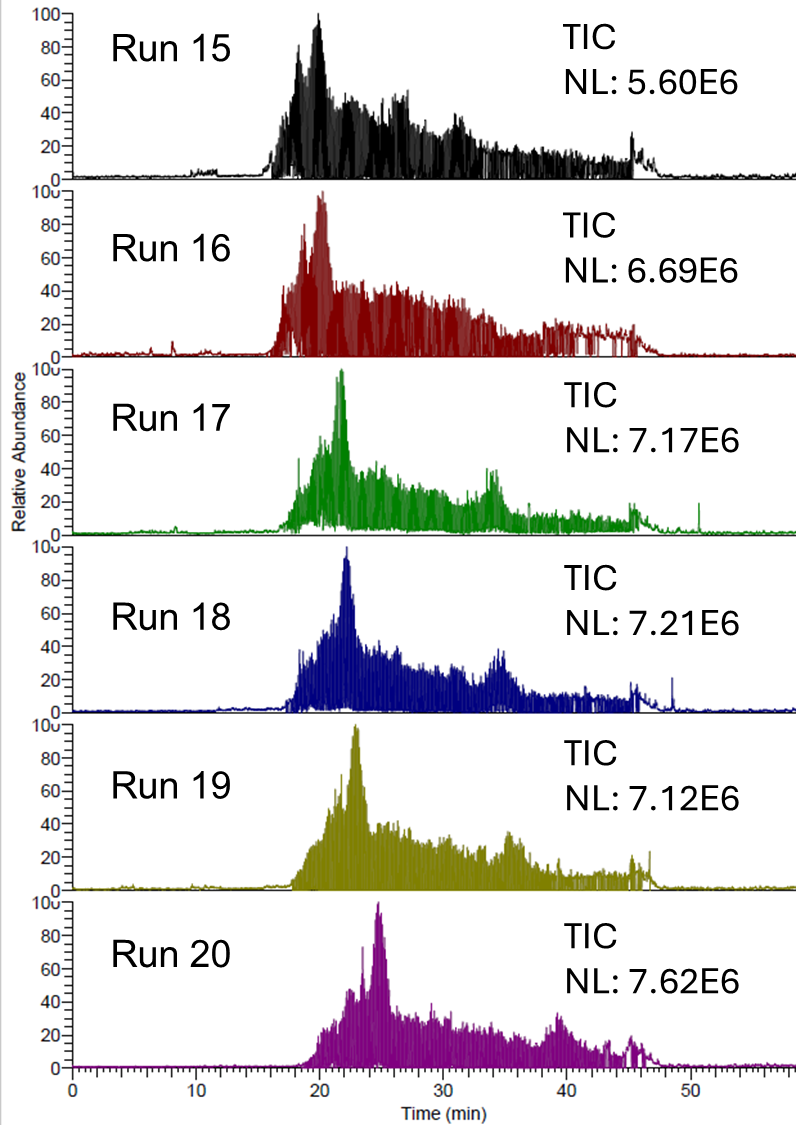

**Figure S13**. Total ion current (TIC) electropherograms of one size exclusion chromatography (SEC) fraction of the HeLa cell lysate after six native CZE-MS/MS runs. The MS1 spectra were acquired under a low-mass-resolution condition (6250, m/z 400), and MS/MS spectra were acquired under a low-mass-resolution condition (6250, m/z 400).

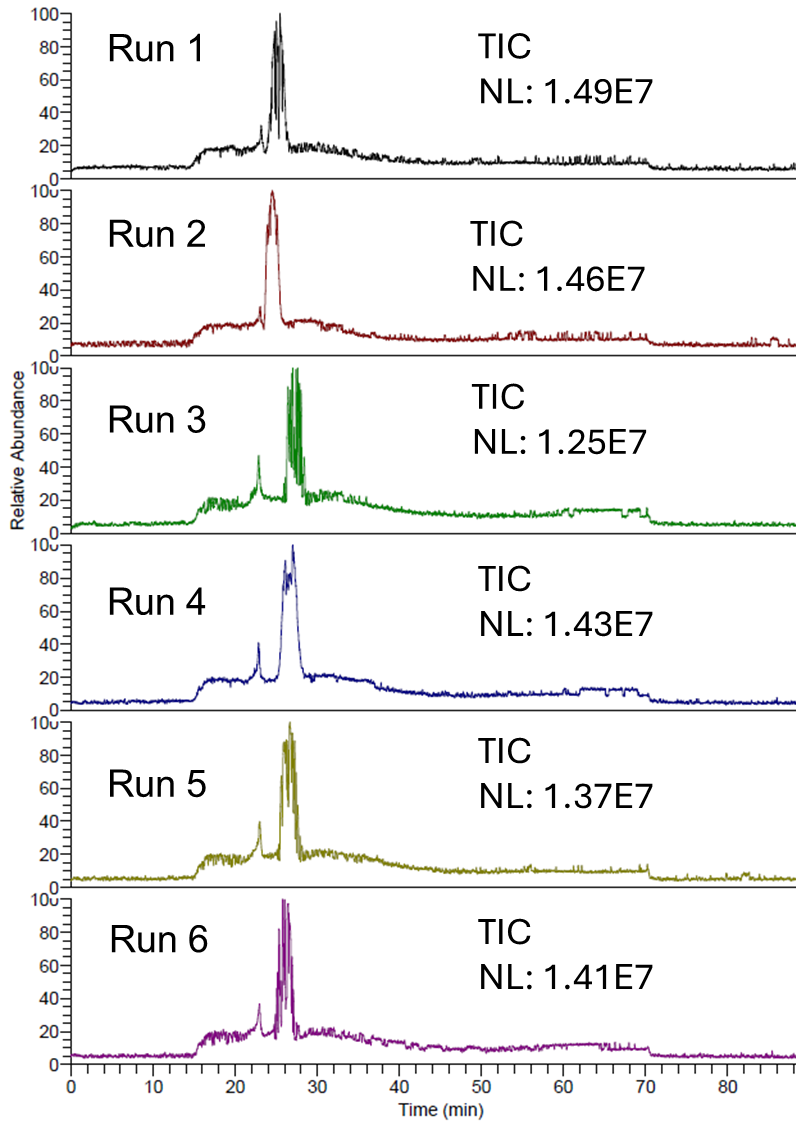

**Figure S14**. Total ion current (TIC) electropherograms of six CZE-MS runs of a human plasma sample. Only MS1 spectra were acquired.

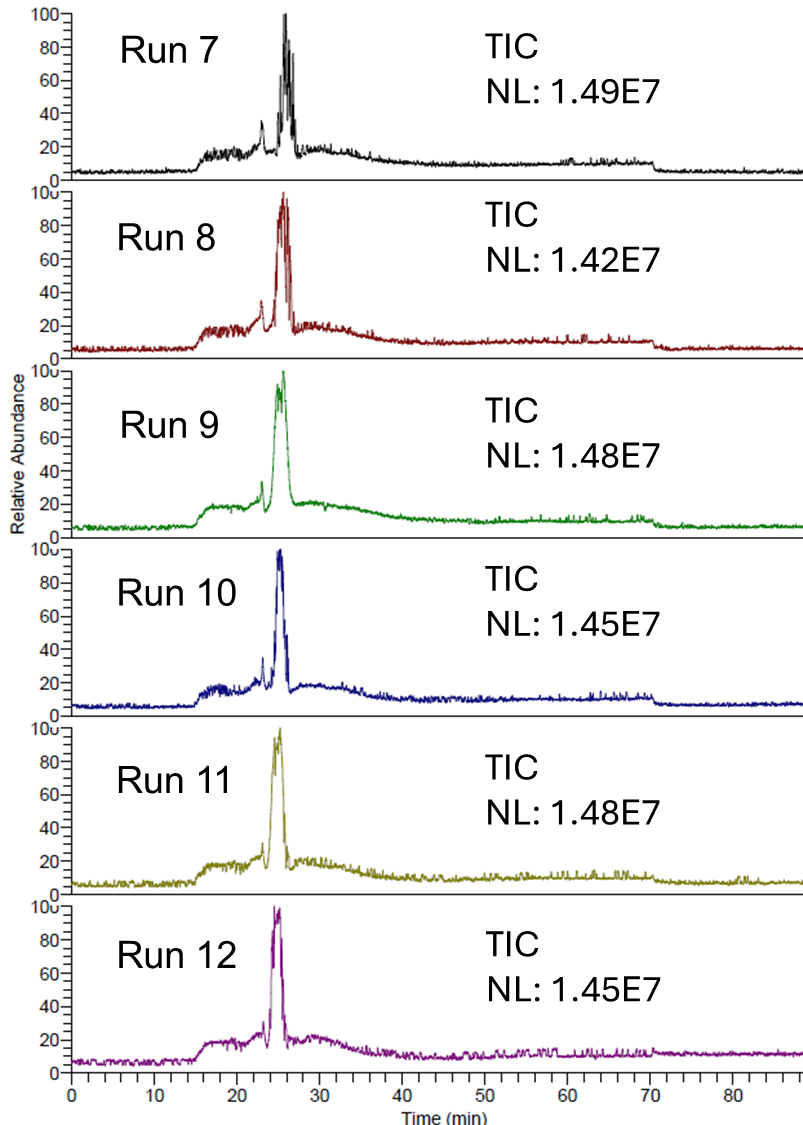

**Figure S15**. Total ion current (TIC) electropherograms of another six CZE-MS runs of a human plasma sample. Only MS1 spectra were acquired.

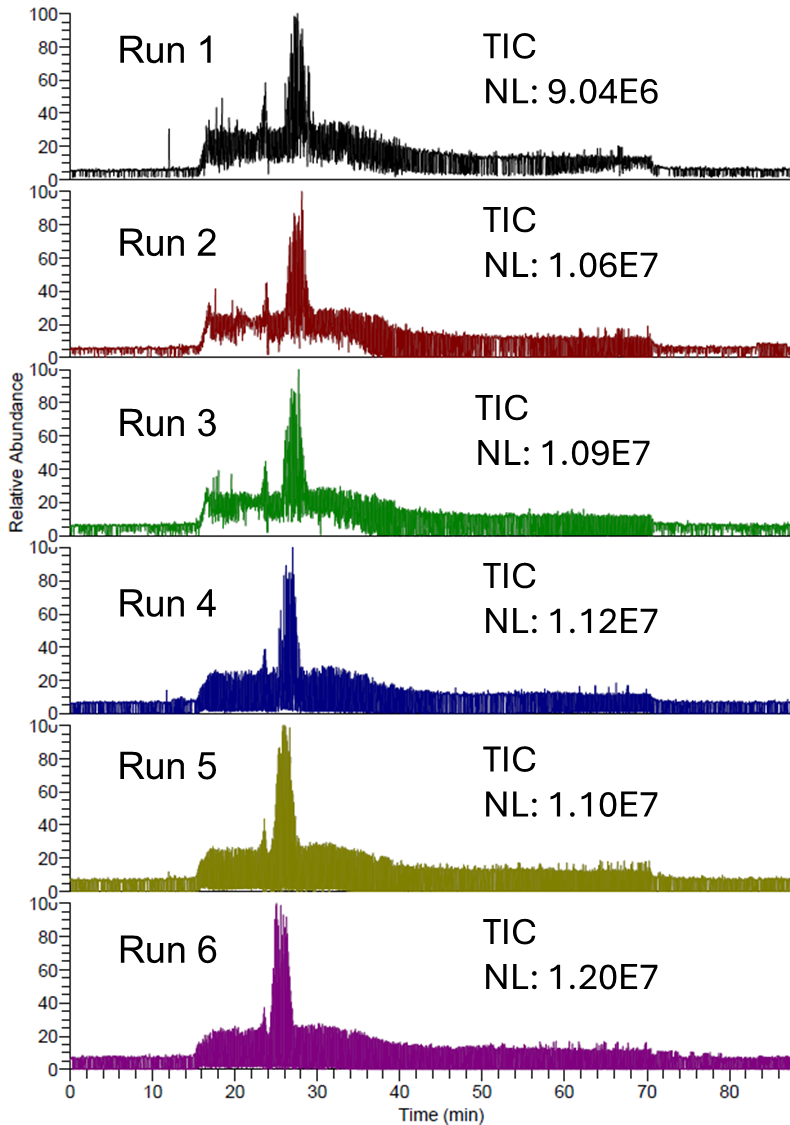

**Figure S16**. Total ion current (TIC) electropherograms of human plasma after six native CZE-MS/MS runs. The MS1 spectra were acquired under a low-mass-resolution condition (6250 at m/z 400). For runs 1, 2, and 3, the MS/MS spectra were acquired under a high-mass-resolution condition (100,000 at m/z 400). For runs 4, 5, and 6, the MS/MS spectra were acquired under a low-mass-resolution condition (6250 at m/z 400).

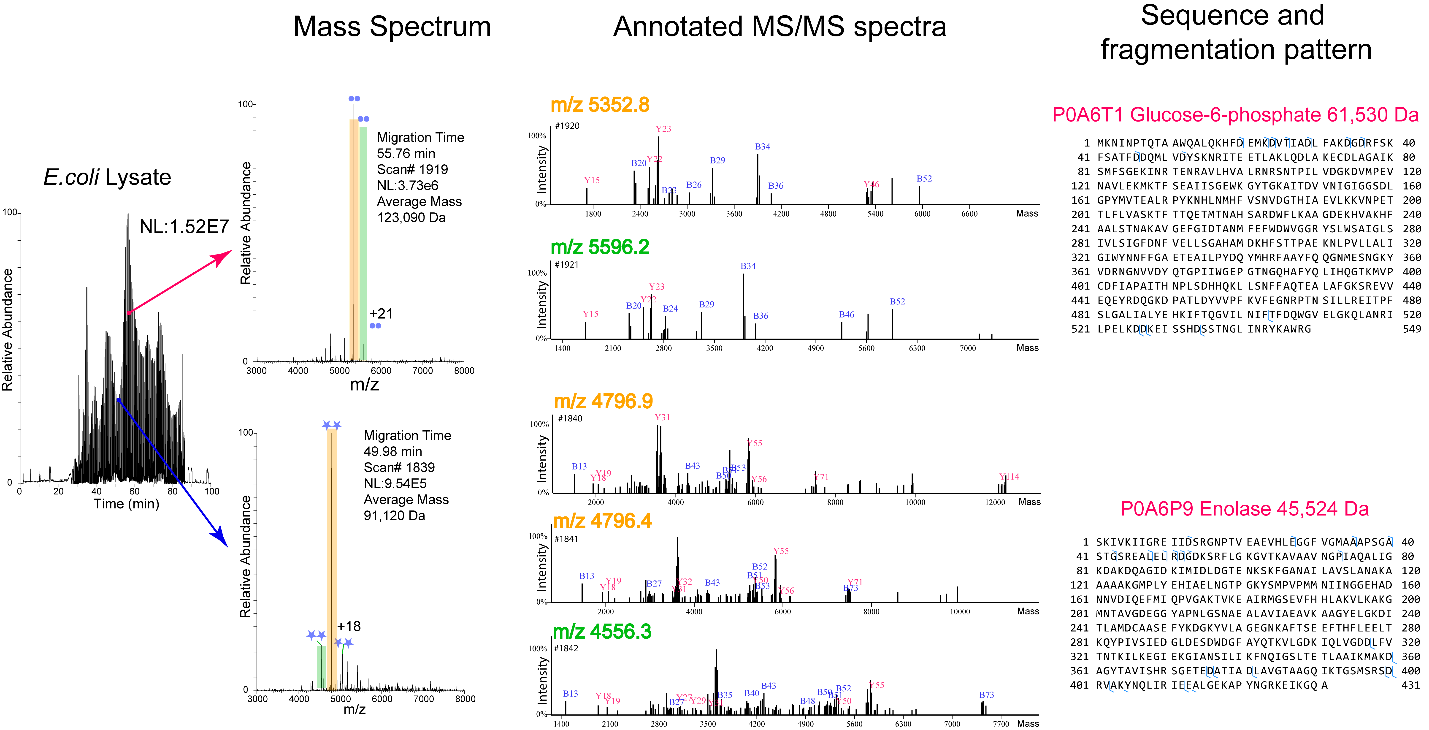

**Figure S17**. Two examples of identified complexoforms from the *E. coli* cell lysate. The TIC electropherogram of one CZE-MS/MS run, mass spectra, and annotated MS/MS spectra of the complexoforms (two dimers), as well as the sequences and fragmentation patterns of corresponding proteins.

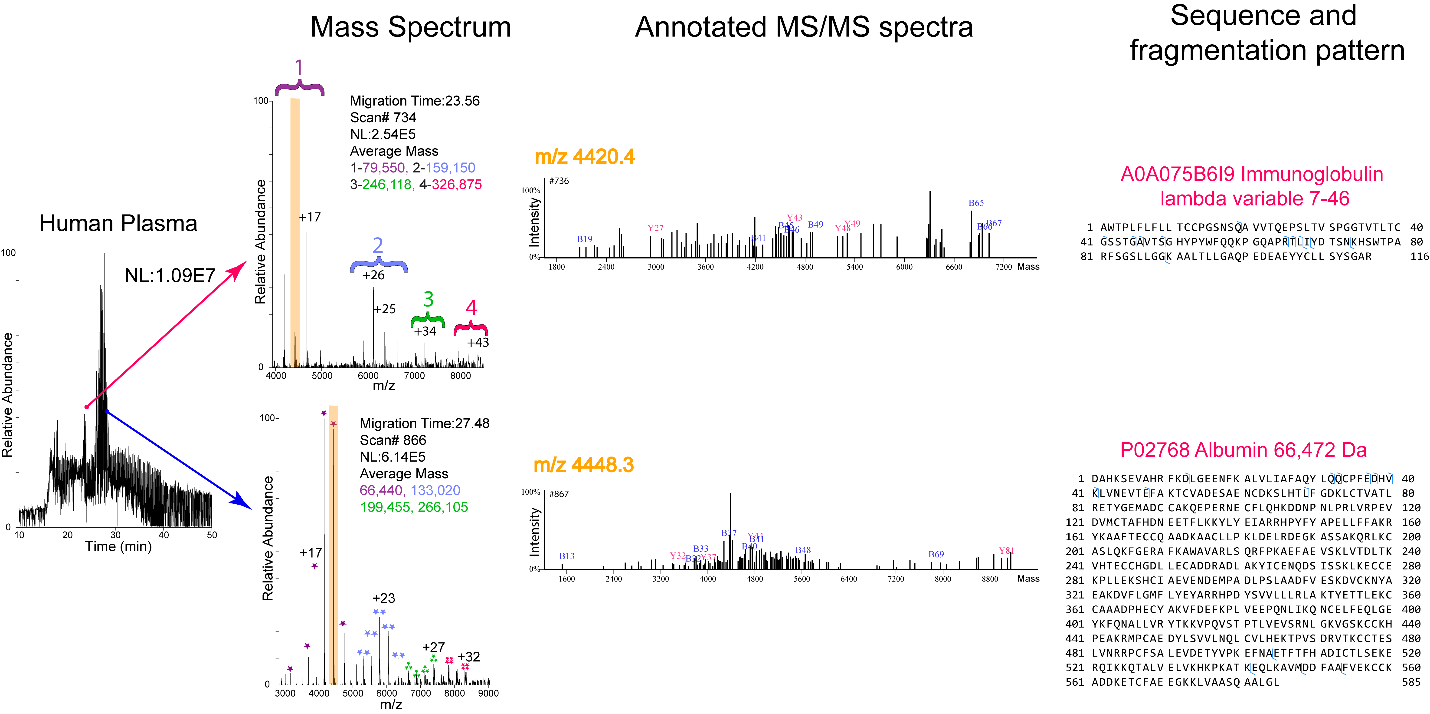

**Figure S18**. Examples of identified complexoforms from the human plasma sample. The TIC electropherogram of one CZE-MS/MS run, mass spectra, and annotated MS/MS spectra of the complexoforms, as well as the sequences and fragmentation patterns of proteins related to the complexoforms.
